# An Unexpected Role of Neutrophils in Clearing Apoptotic Hepatocytes *In Vivo*

**DOI:** 10.1101/2023.02.08.527616

**Authors:** Luyang Cao, Lixiang Ma, Juan Zhao, Xiangyu Wang, Xinzou Fan, Wei Li, Yawen Qi, Yingkui Tang, Jieya Liu, Shengxian Peng, Li Yang, Liangxue Zhou, Li Li, Xiaobo Hu, Yuan Ji, Yingyong Hou, Yi Zhao, Xianming Zhang, Youyang Zhao, Yong Zhao, Yuquan Wei, Asrar B. Malik, Hexige Saiyin, Jingsong Xu

**Author notes:** Correspondence should be addressed to: Jingsong Xu, or Hexige Saiyin. These authors contributed equally to this work (Luyang Cao, Lixiang Ma, Juan Zhao, Xiangyu Wang). Department of Pharmacology, Center for Lung and Vascular Biology, University of Illinois, Chicago (Jingsong Xu).

## Abstract

Billions of apoptotic cells are removed daily in a human adult by professional phagocytes (e.g. macrophages) and neighboring nonprofessional phagocytes (e.g. stromal cells). Despite being a type of professional phagocyte, neutrophils are thought to be excluded from apoptotic sites to avoid tissue inflammation. Here we report a fundamental and unexpected role of neutrophils as the predominant phagocyte responsible for the clearance of apoptotic hepatic cells in the steady state. In contrast to the engulfment of dead cells by macrophages, neutrophils burrowed directly into apoptotic hepatocytes, a process we term *perforocytosis*, and ingested the effete cells from the inside. The depletion of neutrophils caused defective removal of apoptotic bodies, induced tissue injury in the mouse liver and led to the generation of autoantibodies. Human autoimmune liver disease showed similar defects in the neutrophil-mediated clearance of apoptotic hepatic cells. Hence, neutrophils possess a specialized immunologically silent mechanism for the clearance of apoptotic hepatocytes through perforocytosis, and defects in this key housekeeping function of neutrophils contribute to the genesis of autoimmune liver disease.

## Introduction

Apoptosis is a process of programmed cell death that clears aged or damaged cells to maintain internal tissue homeostasis(Hochreiter-Hufford and Ravichandran, 2013). An estimated hundred billion cells undergo apoptosis daily in a human adult(Fond and Ravichandran, 2016). These apoptotic cells must be disposed of promptly and efficiently without littering cytoplasm or causing inflammation(Franz et al., 2006). Defects in the clearance of apoptotic bodies are often linked to various inflammatory and autoimmune diseases(Nagata et al., 2010; Poon et al., 2014). It is widely believed that apoptotic bodies are cleared by professional phagocytes, such as macrophages and immature dendritic cells, or by local nonprofessional phagocytes, such as epithelial cells, endothelial cells, and fibroblasts(Lauber et al., 2004). Although the morphology of apoptotic cells is easily distinguished in histological sections(Kerr et al., 1972), these cells are infrequently observed in normal human samples due to efficient removal by phagocytes(Lauber et al., 2004; Poon et al., 2014). Hence, the phagocytes responsible for the removal of apoptotic cells in the homeostatic state remain uncertain and it is unknown whether they are tissue specific.

Neutrophils, a type of terminally differentiated and short-living phagocytic cell, represent 50-70% of the total white blood cell population in humans(Amulic et al., 2012; Kolaczkowska and Kubes, 2013). However, neutrophils are not considered key players in apoptotic cell clearance(Nagata et al., 2010; Poon et al., 2014). They instead function as inflammatory cells responsible for killing bacteria and fighting infection. Blood neutrophils often swarm to the site of infection, where they release toxic mediators (e.g., reactive oxygen species and proteases) that not only kill microorganisms but also can damage tissues(Amulic et al., 2012; Kolaczkowska and Kubes, 2013). To prevent inflammation, apoptotic cells have been shown to release ‘keep-out’ signals, such as lactoferrin, that prevent the recruitment of neutrophils to the apoptotic site(Bournazou et al., 2009). Nevertheless, neutrophils are related to multiple autoimmune diseases(Amulic et al., 2012; Kolaczkowska and Kubes, 2013). For example, mild neutropenia was observed to precede and accompany the onset of type 1 diabetes, a typical autoimmune disease(Harsunen et al., 2013; Valle et al., 2013). The role of neutrophils in the clearance of apoptotic bodies and contribute to autoimmunity remain to be solved (Jorch and Kubes, 2017).

Here we report an unexpected role of neutrophils in clearing apoptotic hepatocytes under physiological conditions. We found that neutrophils, not macrophages or other cells, are responsible for apoptotic hepatocyte clearance in an immunologically silent manner. We noted that neutrophils penetrate apoptotic hepatocytes and clear them from the inside, thereby avoiding spillage of cytoplasmic content; this mechanism differs sharply from the classical engulfment of dead cells by macrophages. Defects in neutrophil-mediated apoptotic clearance have been observed in human autoimmune liver (AIL) disease. Hence, in addition to their well-known role in combating infection, neutrophils can function as housekeepers for apoptotic clearance and thus maintain tissue homeostasis.

## Results

### Neutrophils burrow into apoptotic hepatocytes in human livers

We observed a large number of neutrophils in hepatocytes in human noncancerous liver tissue obtained from patients with hepatocellular carcinoma (Fig. 1A) or from patients with hepatic hemangioma (Fig. 1B). We discovered that the hepatocytes occupied by neutrophils were apoptotic, as evidenced by the condensed chromatin signature (Fig. 1A, panels i-vi, black arrowheads indicate condensed chromatin, white arrowheads point to neutrophils with a characteristic multilobed nucleus). Importantly, apoptotic hepatocytes are rarely observed in the human liver, possibly due to the rapid removal of apoptotic cells by phagocytes. In a total of 281 apoptotic hepatocytes from 32 livers, we observed that each apoptotic hepatocyte was engorged by up to 22 neutrophils (Fig. 1C, Table S1). We also confirmed apoptotic hepatocytes by TUNEL staining or Caspase-3 immunostaining (Fig. 1D). Neutrophils burrowed inside apoptotic hepatocytes were either stained with an antibody against neutrophil elastase (NE), myeloperoxidase (MPO) or recognized by their multilobed nucleus signature (Fig. 1D, S1A). The distances from burrowed neutrophils to the apoptotic hepatocyte border were analyzed by IMARIS software and recorded in Table S2. Other immune cells were rarely associated with apoptotic hepatocytes. Kupffer cells are liver-resident phagocytes that are thought to clear apoptotic bodies(Canbay et al., 2003; Eipel et al., 2007). Upon staining with an antibody against CD68, a marker of Kupffer cells, we observed few Kupffer cells invading or engulfing apoptotic hepatocytes (Fig. 1C, E, 7A and Table S3). Furthermore, we detected few CD11b^+^ or CD45RA^+^ cells associated with the apoptotic sites (Fig. S1B, Table S4, S5).

**Figure 1.**
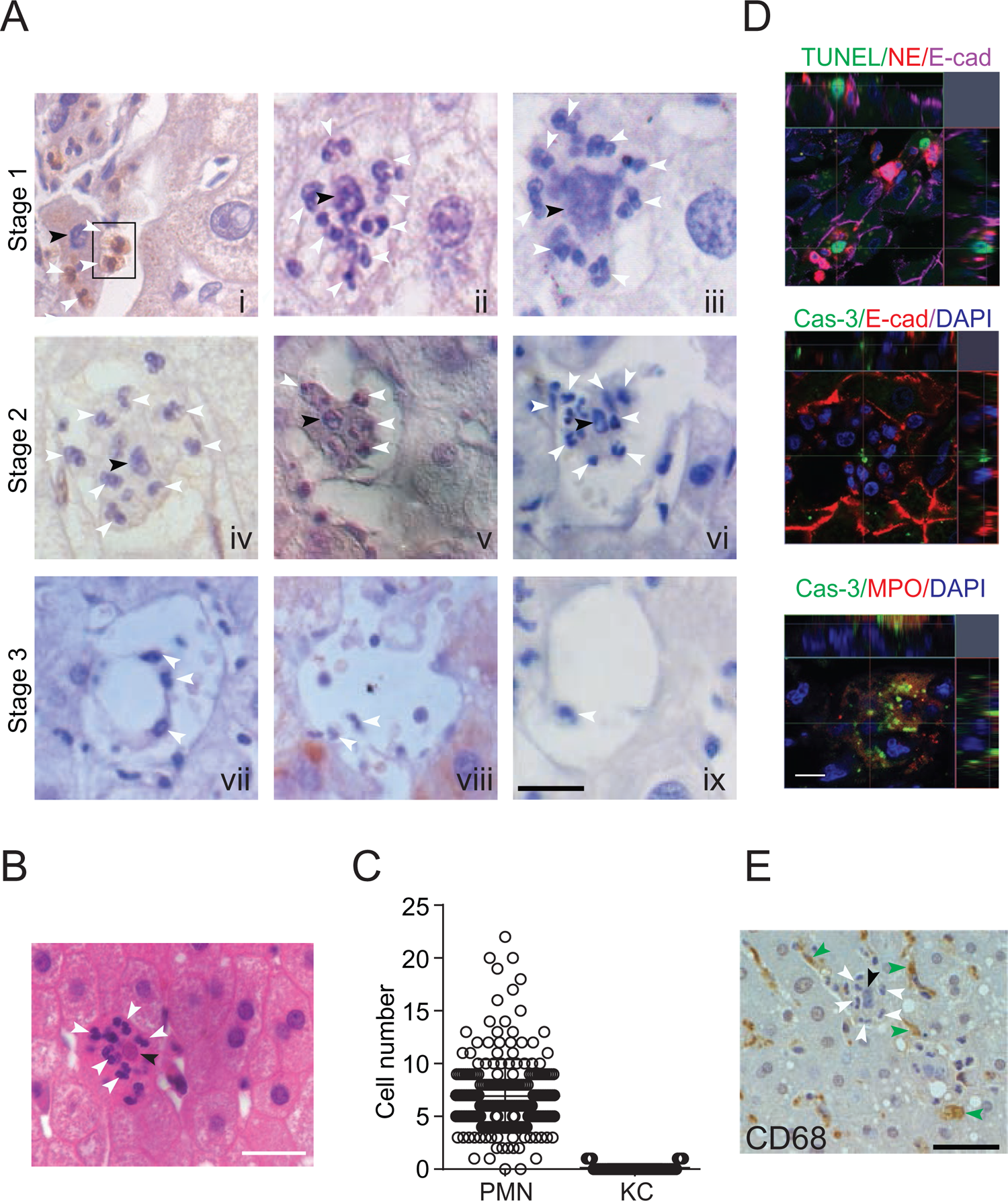
Neutrophil-mediated clearance of apoptotic cells in livers. (A) Hematoxylin staining of human noncancerous liver tissues from patients with hepatocellular carcinoma. Apoptotic hepatocytes with apparent condensed chromatin are being scavenged by several neutrophils (black arrowhead indicates condensed chromatin and white arrowheads point to neutrophils with a characteristic multilobed nucleus). There are three stages of neutrophil scavenging apoptotic hepatocytes in human samples. Stage 1 (i-iii), initial invading stage: two neutrophils start to invade an apoptotic hepatocyte with apparent condensed chromatin (outlined by the black rectangle); three other neutrophils are already inside the cytoplasm (i); at this stage, the apoptotic hepatocytes are still attached to neighboring hepatocytes and occupied by 2-16 neutrophils (ii-iii, 37 out of 241 apoptotic hepatocytes observed are at stage 1). Stage 2, phagocytosis and shrinking stage: The apoptotic hepatocytes are being phagocytosed by invading neutrophils and are partially detached from neighboring hepatocytes (iv-vi, 130 out of 241 apoptotic cells are at stage 2). Stage 3, complete digestion stage: apoptotic hepatocytes are completely detached and largely cleared, and only neutrophils remain in the cleared region (vii-ix, 74 out of 241 apoptotic cells are at stage 3). Scale bar, 20 μm. (B) Hematoxylin staining of human liver tissues from patients with hemangioma (a total of 40 apoptotic hepatocytes were observed). Scale bar, 20 μm. (C) Cell counts of neutrophils (PMNs) and Kupffer cells (KCs) inside or associated with apoptotic hepatocytes, also see Table S1 and S3. (D) Fluorescent images of human liver tissues with neutrophils scavenging apoptotic hepatocytes. Apoptotic hepatocytes are confirmed with TUNEL staining or Caspase-3 immunostaining. Neutrophils are labelled with immunostaining of neutrophil elastase (NE), myeloperoxidase (MPO) or DAPI staining (with a segmented nucleus signature). Scale bar, 10 μm. (E) Images of liver tissue stained with an antibody against CD68, a marker of KCs (indicated by green arrowheads). KCs do not invade or phagocytose apoptotic hepatocytes (black arrowheads). Data are representative of (A, B, D, E) or from (C) a total of 281 apoptotic hepatocytes observed, mean and s.e.m. in C.

Based on the above observations that neutrophils are the predominant phagocyte associated with apoptotic hepatocytes, we hypothesized that the neutrophil-mediated clearance of apoptotic cells consists of the following three sequential steps. In the initial invading or burrowing stage, activated neutrophils identified and targeted the apoptotic hepatocytes (cells with condensed chromatin, indicated by black arrowheads) and attached to their cell membrane (Fig. 1A, panel i, outlined by black rectangle). Then, the neutrophils invaded apoptotic hepatocytes (Fig. 1A, panels ii and iii). We observed an average of 7 neutrophils entering each apoptotic hepatocyte, and we termed this process perforocytosis (from Latin *perfero* to bore) (Fig. 1A, white arrowheads point to neutrophils). The second step consisted of phagocytosis and detachment. The neutrophils within hepatocytes appeared to clear apoptotic bodies from the inside without destroying the cellular membrane or extruding the cytoplasm (two burrowed neutrophils phagocytosed apoptotic debris as shown in Fig. S1C). Following digestion by neutrophils, the apoptotic hepatocytes decreased in size, and detached from nearby hepatocytes (Fig. 1A, panels iv-vi). The third step involved the complete digestion of apoptotic hepatocytes (Fig. 1A, panels vii-ix). After the clearance of apoptotic hepatocytes, neutrophils seemed to migrate away from the cleared space, possibly to make room for new hepatocytes generated by rapid division. This ‘eating inside’ phagocytosis process differed sharply from the well-known engulfment of apoptotic cells or fragmented apoptotic bodies by other phagocytic cells, such as macrophages(Poon et al., 2014).

### Visualization of neutrophil perforocytosis in mouse livers

Neutrophils within apoptotic hepatocytes were confirmed in mouse livers by intravital microscopy (Fig. 2A, B) and electron microscopy (Fig. 2C). Similar to the observations in human samples, apoptotic hepatocytes in WT mouse livers were occupied by neutrophils but not associated with macrophages (Fig. 2A, neutrophils were labeled with an i.v. injection of anti-Ly6G antibody, macrophages were labeled with anti-F4/80 antibody and apoptotic cells were labeled with Annexin V). A total of 24 apoptotic cells were observed in 8 WT livers with an average of 2 burrowed neutrophils in each mouse apoptotic hepatocyte. The burrowed neutrophils were projected and analyzed by IMARIS software and results were compared side by side with livers from MRP8cre/DTR mice (Fig. 2A, with more details in the neutrophil depletion section below). The distances of burrowed neutrophils to the border of apoptotic hepatocytes were recorded in Table S6.

**Figure 2.**
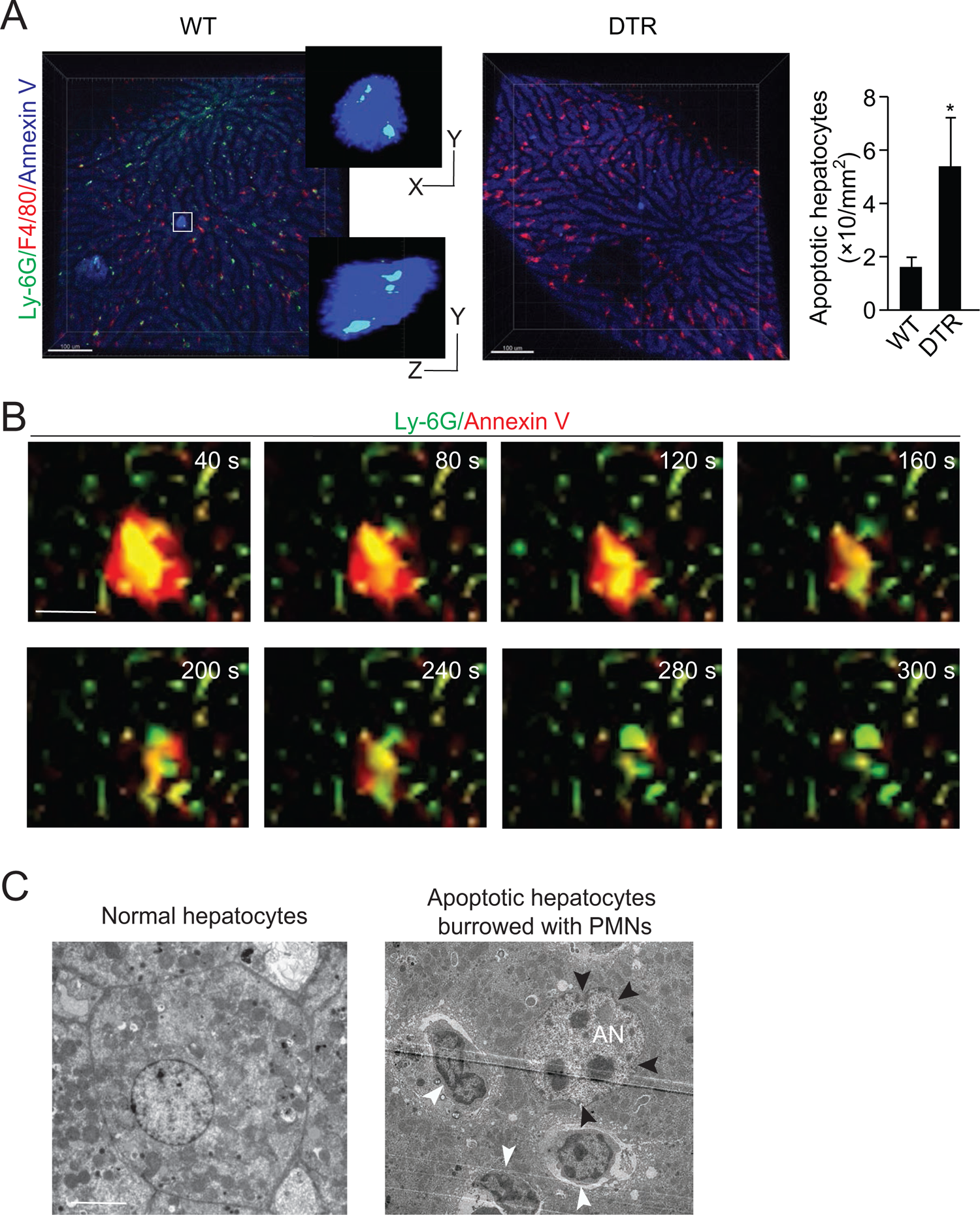
Neutrophil burrowing into apoptotic hepatocytes. (A) Intravital microscopy images of liver tissues from WT and MRP8cre/DTR mice. Neutrophils are labeled with an i.v. injection of anti-Ly6G antibody (green), KCs are labeled with anti-F4/80 antibody (red), and apoptotic cells are labeled with Annexin V (blue). Neutrophils inside and KCs associated with apoptotic hepatocytes are detected and analyzed with IMARIS software as described in Methods. The distances from neutrophils to the apoptotic hepatocyte border are recorded in table S6. A total of 24 apoptotic cells were observed in the WT liver with an average of 2 burrowed neutrophils. Scale bar, 100 μm. (B) Intravital image sequences of neutrophils phagocytosing apoptotic hepatocytes in mouse livers at indicated time points. Neutrophils are labeled with an i.v. injection of anti-Ly6G antibody (green), and apoptotic cells are labeled with Annexin V (red). A total of 13 apoptotic cells with burrowed neutrophils were observed in 12 WT mouse livers. Scale bar, 20 μm. (C) Electron microscopy images of apoptotic mouse hepatocytes occupied by neutrophils. The apoptotic hepatic nucleus (AN) is evident by distorted nuclear membrane (pointed by black arrowheads). The neutrophils are indicated by white arrowheads with a characteristic multilobed nucleus. 29 apoptotic cells with burrowed neutrophils were observed. Scale bar, 5 μm. Data are representative of three independent experiments.

By using the Cellvizio System (Confocal Miniprobes), we managed to visualize the entire process of neutrophil perforocytosis in mouse livers (Fig. 2B and Video S1, S2, apoptotic hepatocytes were labeled with Annexin V and neutrophils were labeled with anti-Ly6G antibody). Consistent with observations in human samples, Annexin V positive apoptotic hepatocytes were burrowed and cleared by Ly6G-labeled neutrophils (Fig. 2B). This eating inside phagocytic process is fast and rigorous in which neutrophils were able to completely digest apoptotic hepatocytes around 4-7 minutes (Fig. 2B, and Video S1, S2, a total of 13 apoptotic hepatocytes in 12 WT mouse livers were observed). At the end of the apoptotic clearance process, neutrophils simply left the apoptotic sites and were not labeled by Annexin V, indicating neutrophils were not apoptotic.

### The selectivity of neutrophil perforocytosis of effete hepatocytes

To study neutrophil-mediated phagocytosis in live cells in vitro, we induced apoptosis of isolated human primary hepatocytes or mouse liver NCTC cells with puromycin (Fig. S2A) and then applied human neutrophils or neutrophil-like, differentiated HL60 cells (Fig. 3). Isolated human primary hepatocytes formed a hepatic plate-like structure around 7 days in the culture dishes and were further confirmed with anti-albumin antibody (Fig. 3A). Human blood neutrophils (labeled with a red membrane dye, PKH-26) were able to penetrate and burrow inside the human apoptotic hepatocytes induced by puromycin (Fig. 3B, white arrowheads point to burrowing neutrophils). These apoptotic hepatocytes had markedly decrease in size after neutrophil burrowing as compared with cells without neutrophil burrowing (Fig. 3C, D, Video S3 shows a human neutrophil first burrowed and then started to phagocytose an apoptotic hepatocyte from inside in real time). Similar results were observed with NCTC cells (labeled with a red membrane dye, PKH-26) and HL60 cells (labeled with a green membrane dye, PKH-67). Green-labeled HL60 cells showed little response to normal NCTC cells (Fig. 3E, top row). In the presence of apoptotic NCTC cells, however, green-labeled HL60 cells polarized and penetrated dead cells (Fig. 3E, white arrowheads point to burrowing neutrophils, bottom row). Next, we quantified phagocytosis by flow cytometry using a pH sensitive dye, PHrodo (with weak fluorescent at neutral pH but high fluorescent with a drop in pH to measure engulfment, see Methods). The phagocytosis of apoptotic hepatocytes by neutrophils was markedly greater than that of normal nonapoptotic hepatocytes (Fig. 3F, G). There was little cytoplasm or DNA leakage during neutrophils phagocytosing apoptotic NCTC cells as assessed by the extracellular levels of DNA, SOD and ROS (Fig. S1D-F). To address whether neutrophils also phagocytose other apoptotic cells, we determined the ability of neutrophils to phagocytose apoptotic endothelial HUVECs or epithelial HEK293 cells. We did not observe phagocytosis of these cells compared with their nonapoptotic controls (Fig. 3G), indicating the selectivity of neutrophils in phagocytosing apoptotic hepatocytes. Upon comparing the ability of HL60 cells and macrophage U937 cells to phagocytose apoptotic hepatocytes, we found that U937 cells exhibited much lower phagocytosis of apoptotic NCTC cells than did HL60 cells (Fig. S2C). Similar results were observed with human and mouse macrophages (Fig. S2B, C), consistent with the central role of neutrophils in mediating apoptotic hepatocyte clearance.

**Figure 3.**
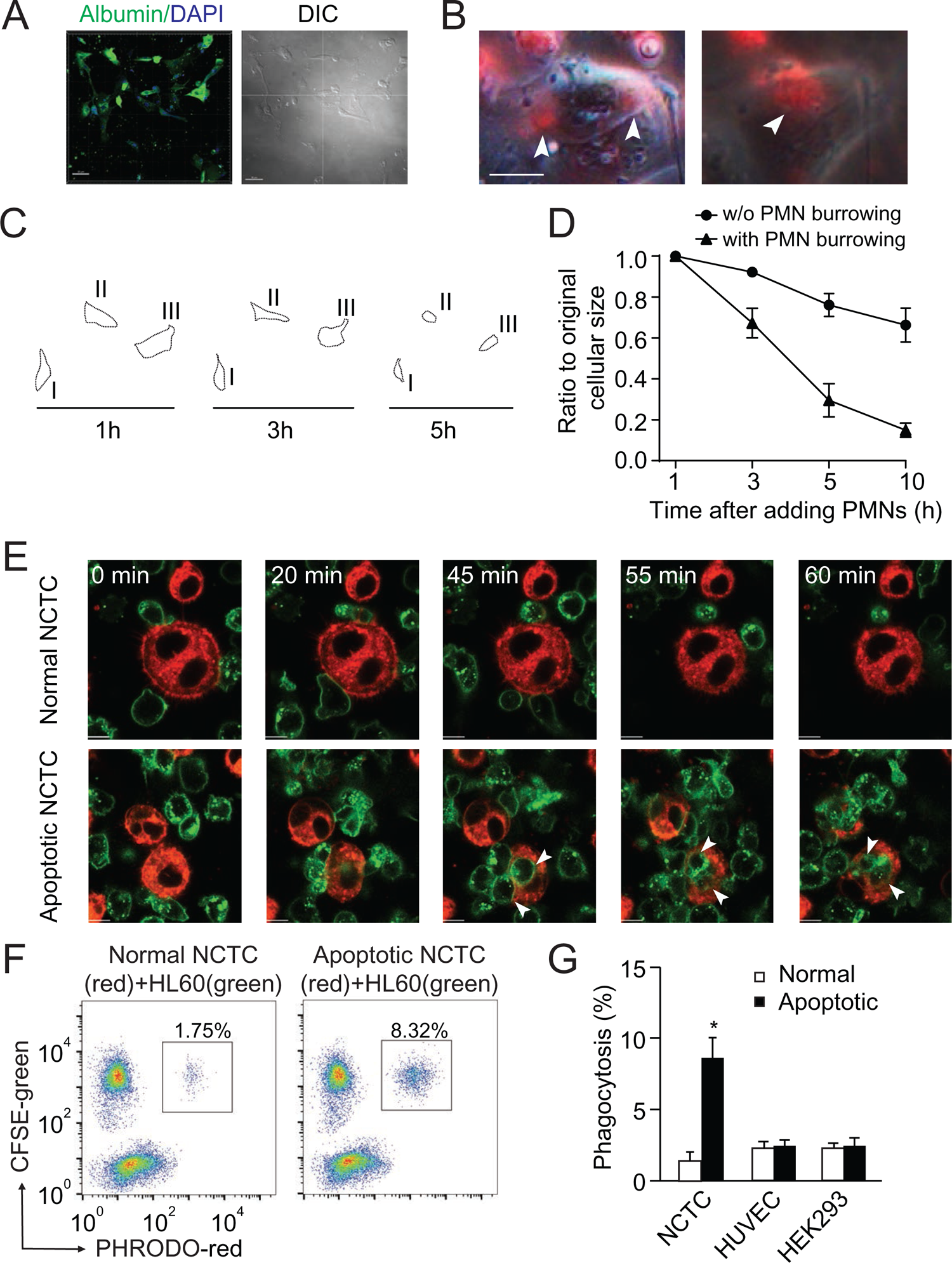
Neutrophils preferentially phagocytose apoptotic hepatocytes. (A) Fluorescent and phase contract images of isolated and cultured human primary hepatocytes stained with anti-albumin antibody (green). Scale bar, 50μm. (B) Fluorescent and phase contract images of burrowed human neutrophils (red) inside apoptotic human primary hepatocytes from A (treated with puromycin). Scale bar, 10μm. (C, D) Cell outlines (C) and cell size quantification (D) of apoptotic human primary hepatocytes occupied with burrowed neutrophils at indicated time points. (E) Fluorescence images of PKH67 (green)-labeled HL60 cells interacting with PKH26 (red)-labeled NCTC cells at the indicated time points. HL60 cells exhibited littler responses to nonapoptotic NCTC cells (top row). In contrast, HL60 cells polarized and penetrated apoptotic NCTC cells (bottom row). Scale bar, 10 μm. (F) Flow cytometry analysis of HL60 cells (labeled with CFSE, green) incubated with nonapoptotic (first column) or apoptotic hepatocytes (second column, hepatocytes were labeled with PHrodo-red dye with weak fluorescent at neutral pH but high fluorescent with a drop in pH). The population of double fluorescent HL60 cells in this subgroup (with higher PHrodo-red fluorescent indicating engulfment) was calculated, and these cells were considered phagocytosing cells. (G) Quantification of F. HL60 cells exhibited significantly higher phagocytosis of apoptotic NCTC cells than that of nonapoptotic controls. *, *P* < 0.05 (Student’s *t*-test). Data are representative of (A-C, E, F) or from (D, G) three independent experiments, mean and s.e.m. in D, G.

### The signals that attract neutrophils to apoptotic hepatocytes

To identify the signals that attract neutrophils to apoptotic hepatocytes, we screened for cytokine and chemokine secreted by NCTC cells before and after apoptosis. Apoptotic NCTC cells showed markedly increased secretion of the cytokines IL-1β, IL-6, IL-8, and IL-12 compared with nonapoptotic controls (Fig. 4A-D). In contrast, GM-CSF, IFN-γ, TNF-α, IL-2 and IL-10 were not significantly changed (Fig. S3A). To address the role of the upregulated cytokines, we knocked down the receptors of IL-1β, IL-6, IL-8, or IL-12 in HL60 cells and then examined phagocytosis ability. Knockdown of the IL-1β receptor abolished the phagocytosis of apoptotic NCTC cells (Fig. 4E), whereas knockdown of the IL-8 receptor showed a 50% reduction in phagocytosis (Fig. 4E). Thus, neutrophil chemoattractants, IL-1β and IL-8 secreted by apoptotic NCTC cells are key signals for attracting neutrophils and inducing subsequent phagocytosis. Consistent with above observations, we found endogenous IL-1β associated with apoptotic hepatocytes but not with normal hepatocytes in human liver samples (Fig. 4F).

**Figure 4.**
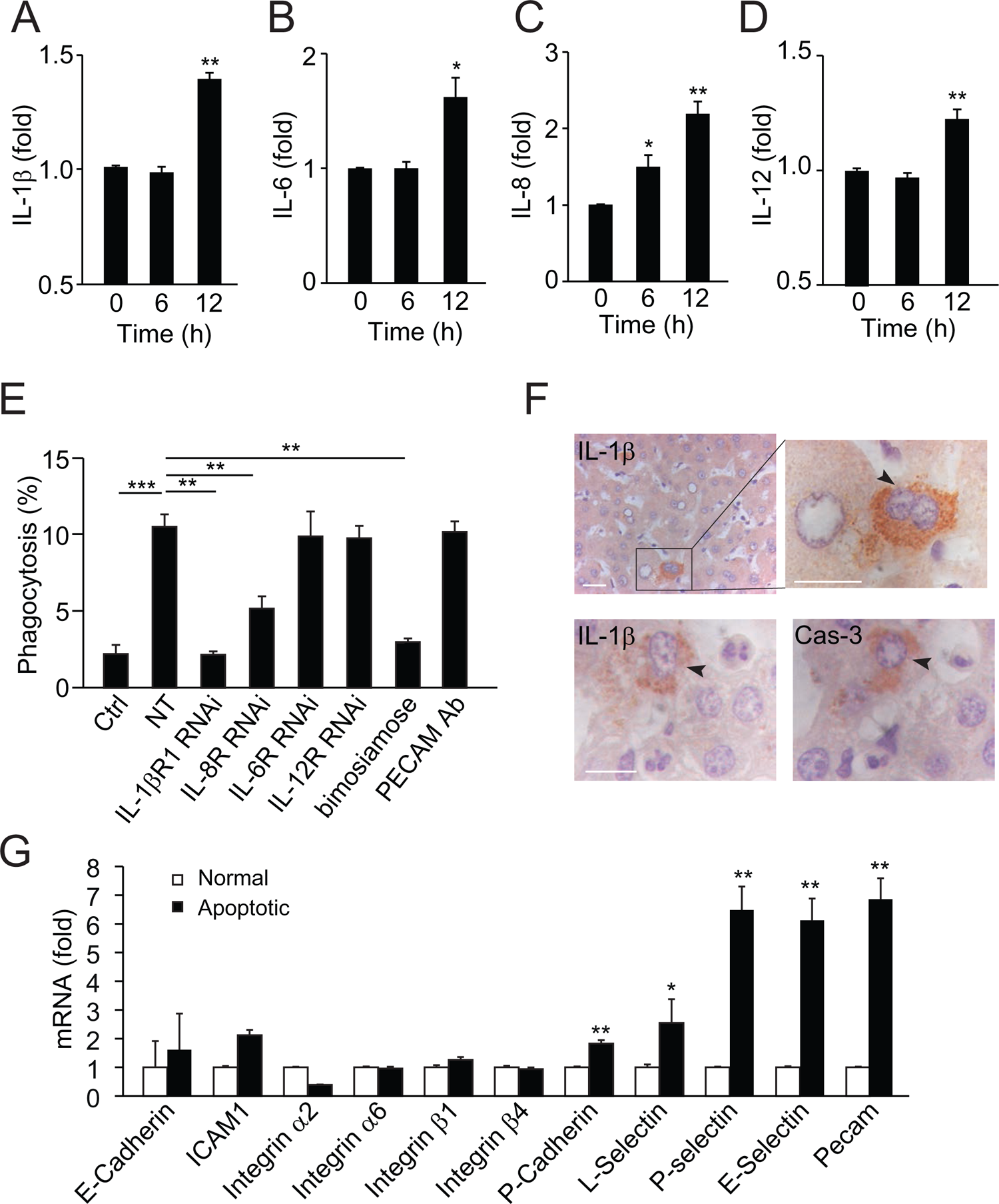
Apoptotic hepatocytes release signals that attract neutrophils for phagocytosis. (A-D) Cytokine secretion in apoptotic NCTC cells. IL-1β (A), IL-6 (B), IL-8 (C), and IL-12 (D) levels were significantly increased in apoptotic cells compared with nonapoptotic controls, **, *P* < 0.01, ***, *P* < 0.001 (Student’s *t*-test). (E) Phagocytosis of nonapoptotic NCTC cells by HL60 cells (ctrl) and of apoptotic NCTC cells by HL60 cells treated without (NT) or with RNAi knockdown of IL-1β, IL-8, IL-6, IL-12 receptors, or with selectin antagonist (baimosiamose), PECAM blocking antibody. *, *P* < 0.05 (Student’s *t*-test). (F) Images of liver tissues stained with anti-IL-1β or anti-Cas-3. Top row, IL-1β is only associated with apoptotic hepatocytes with condensed chromatin (indicated by black arrowheads) but not with nonapoptotic hepatocytes. Bottom row, IL-1β and Cas-3 staining in 2μm consecutive liver sections. Both IL-1β and Cas-3 signals are accumulated in the same apoptotic hepatocyte (indicated by black arrowhead). Scale bar, 20 μm. (G) Cell surface receptors in apoptotic NCTC cells compared with nonapoptotic controls. **, *P* < 0.01 (Student’s *t*-test). Data are from (A-E, G) or representative of three independent experiments (F), mean and s.e.m. in A-E, G.

Next, we examined the cell surface molecules in hepatocytes and found that apoptotic NCTC cells had markedly increased P-selectin, E-selection, L-selectin and PECAM as compared with non-apoptotic controls (Fig. 4G). Inhibition of selectins (with a pan-selectin antagonist, bimosiamose) but not PECAM (with a function blocking antibody) abrogated the neutrophil-mediated clearance of apoptotic hepatocytes (Fig. 4E), indicating selectins play a critical role during this process.

### Neutrophil depletion impairs the clearance of apoptotic hepatocytes

We next examined whether neutrophil depletion with either an antibody or genetic methods (see Methods) influences the clearance of apoptotic hepatocytes. Antibody or genetic depletion yielded about 90% or 70% reduction in mouse peripheral blood neutrophils, respectively (Fig. S4A, B). We analyzed liver samples from neutrophil-depleted and control mice by intravital microscopy (Fig. 2A, genetic depletion), or immunostaining (Fig. 5, antibody depletion). Similar to human liver samples, apoptotic hepatic cells in control WT mice were occupied and phagocytosed by neutrophils (Fig. 2A, 5A, B). After neutrophil depletion, the apoptotic bodies were no longer associated with neutrophils (Fig. 2A, 5A, B). However, we observed that macrophages were associated with apoptotic hepatocytes in neutrophil-depleted samples (Fig. 2A, 5A, C), suggesting a compensatory role of macrophages in phagocytosing dead hepatocytes in the absence of neutrophils. The percentage of apoptotic hepatocytes in neutrophil-depleted samples was significantly increased compared to that in controls (0.92% VS 0.2%, *p* < 0.001, Fig. 5D). Hence, neutrophil depletion impaired the prompt clearance of apoptotic cells in the mouse liver. To rule out the effects of environmental microbes after neutrophil depletion, we also treated the neutrophil-depleted mice with antibiotic (20 mg/kg ampicillin, i.p. injected daily). We obtained similar results in neutrophil-depleted mice with or without antibiotic treatment (Fig. 5A-D). Meanwhile, we observed impaired liver function in neutrophil-depleted livers (with increased aspartate aminotransferase and alanine aminotransferase activity and increased total or direct bilirubin level) compared with nontreated controls (Fig. 6A-D, antibody depletion). Depletion of neutrophils had little effect on other tissues (e.g. kidney, Fig. S4C, D).

**Figure 5.**
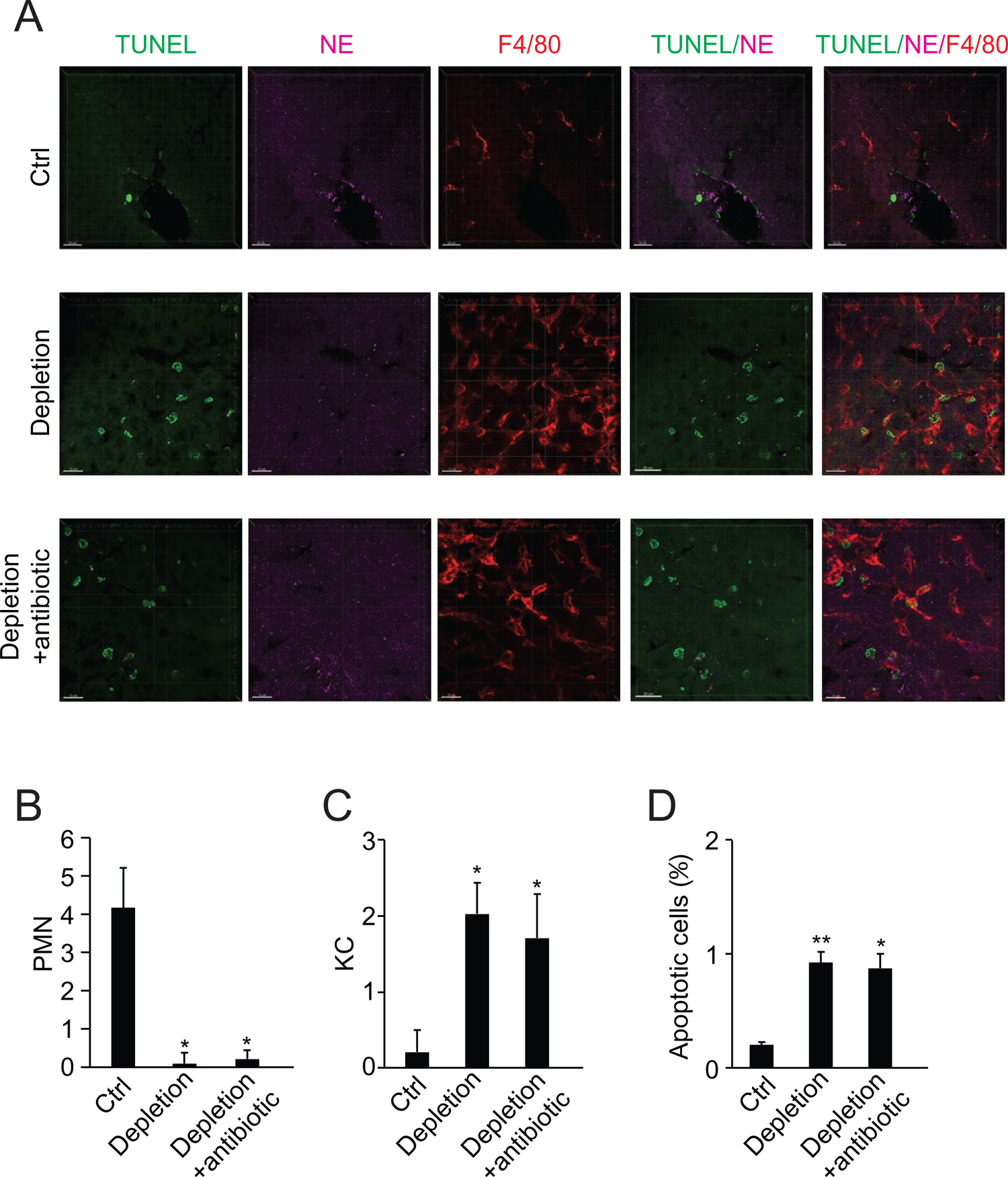
Impaired apoptotic clearance in the neutrophil-depleted liver. (A) Fluorescent images of liver samples from control mice (top row), or neutrophil-depleted mice (antibody depletion; without or with antibiotic, second and third rows). Neutrophils are labeled with NE immunostaining (purple), macrophages are labeled with F4/80 immunostaining (red), and apoptotic cells are labeled with TUNEL staining (green). Scale bar, 15 μm. (B-D) Cell counts of neutrophils (B) and macrophages (C) in or associated with apoptotic hepatocytes (D) in tissue samples as described in a. *, *P* < 0.05, **, *P* < 0.01, compared to control (Student’s *t*-test). Data are representative of (A) or from three independent experiments (B-D; mean and s.e.m. in B-D).

**Figure 6.**
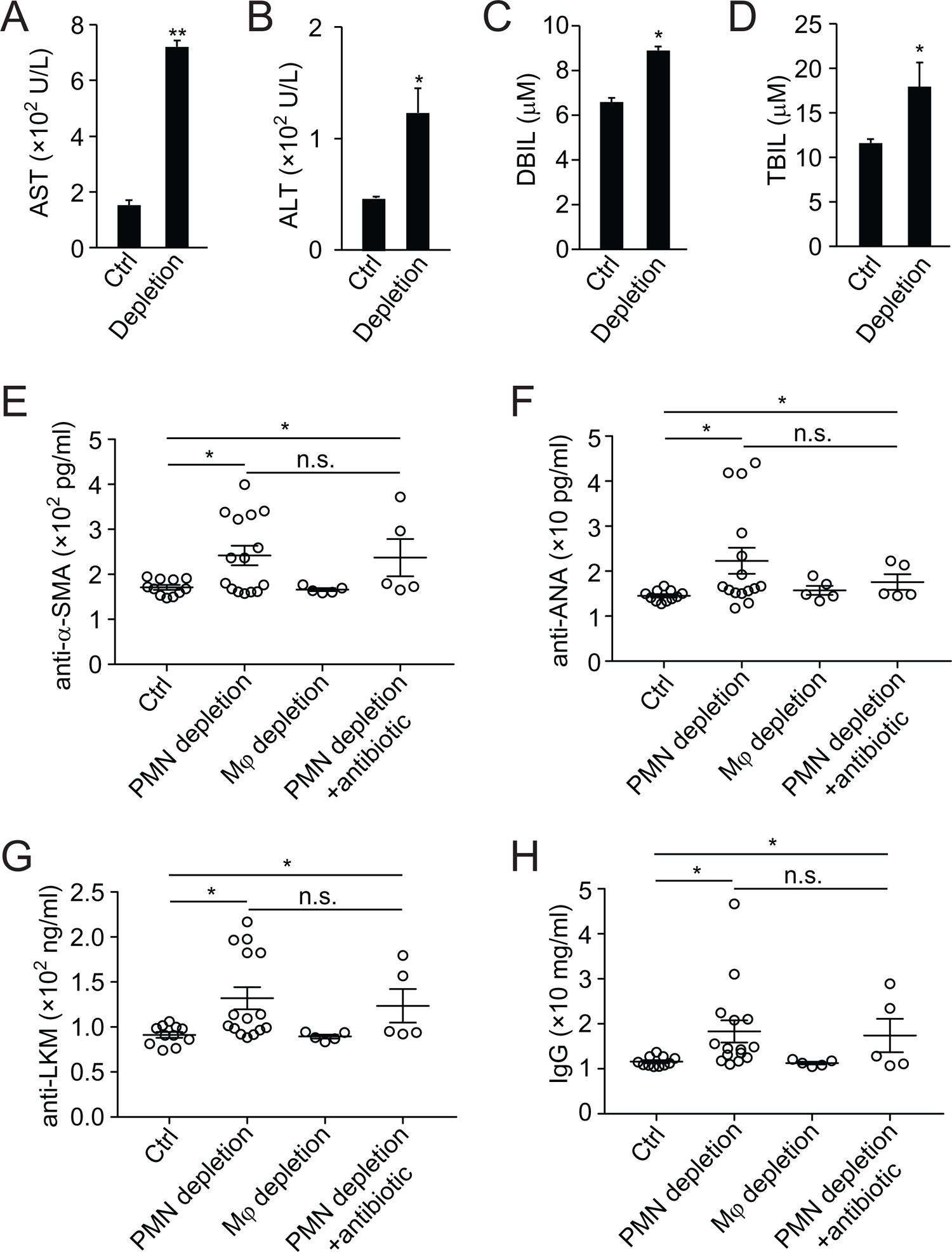
Analysis of liver function and autoantibody generation in neutrophil-depleted mice. (A-D) Liver function analysis of aspartate aminotransferase (AST, A), alanine aminotransferase (ALT, B), direct bilirubin (DBIL, C) and total bilirubin (TBIL, D) in ctrl or neutrophil-depleted mice (antibody depletion). *, *P* < 0.05, **, *P* < 0.01 (Student’s *t*-test). (E-H) Expression of autoantibodies against smooth muscle actin (anti-α-SMA, E), antinuclear antigen (anti-ANA, G), liver-kidney microsome (anti-LKM, G), and total IgG (H) in serum of control, neutrophil-depleted (with antibody), macrophage-depleted, or neutrophil-depleted plus antibiotic treated mice. Data are from three independent experiments (A-H; mean and s.e.m. in A-H).

### Defective neutrophil perforocytosis in autoimmune liver disease

Autoimmune diseases are often linked to defective clearance of apoptotic cells(Nagata et al., 2010; Poon et al., 2014). We first determined whether defective neutrophil-mediated apoptotic clearance contributes to AIL disease. In the present study, we detected an increase in autoantibodies (i.e., against antinuclear antigen, smooth muscle actin, liver-kidney microsome and total IgG antibodies) in neutrophil-depleted mice compared with controls (Fig. 6E-H, antibody depletion). This increase was not affected by the treatment of antibiotic and was not observed in macrophage-depleted mice (Fig. 6E-H, macrophages were depleted with clodronate-liposome, see Methods). Hence, neutrophil depletion not only impaired apoptotic hepatocyte clearance but also led to the generation of autoantibodies, suggesting a role of defective neutrophil-mediated removal of apoptotic bodies in AIL disease.

Next, to address whether neutrophil-mediated apoptotic clearance is impaired in AIL disease, we analyzed biopsy samples from patients diagnosed with AIL disease. In contrast to the normal human controls, a total of 22 AIL disease patient samples contained apoptotic hepatocytes that were not phagocytosed or invaded by neutrophils (Fig. 7A). More apoptotic bodies were associated with macrophages, as observed in neutrophil-depleted mouse livers (Fig. 7A, Table S7). Since the blood neutrophil count in AIL disease patients is within the normal range, we surmised potential defects in the phagocytosis ability of neutrophils from AIL disease patients. We observed markedly decreased phagocytosis of apoptotic NCTC cells by neutrophils from AIL disease patients compared to normal controls (Fig. 7B). Normal human neutrophils burrowed into apoptotic NCTC cells and demonstrated perforocytosis, while AIL disease neutrophils exhibited little response towards apoptotic NCTC cells (Fig. 7C). We further screened differential gene expression between normal human neutrophils and AIL neutrophils. We noted that the expression of IL-1β receptor, IL1R1 and selectin binding protein, P-selectin glycoprotein ligand 1 (PSGL-1) was markedly decreased in AIL neutrophils as compared with normal human neutrophils (Fig. 7D), which further confirmed the critical roles of IL-1β and selectins in neutrophil-mediated apoptotic clearance. The above data prove defective neutrophil-mediated apoptotic clearance in human AIL disease samples.

**Figure 7.**
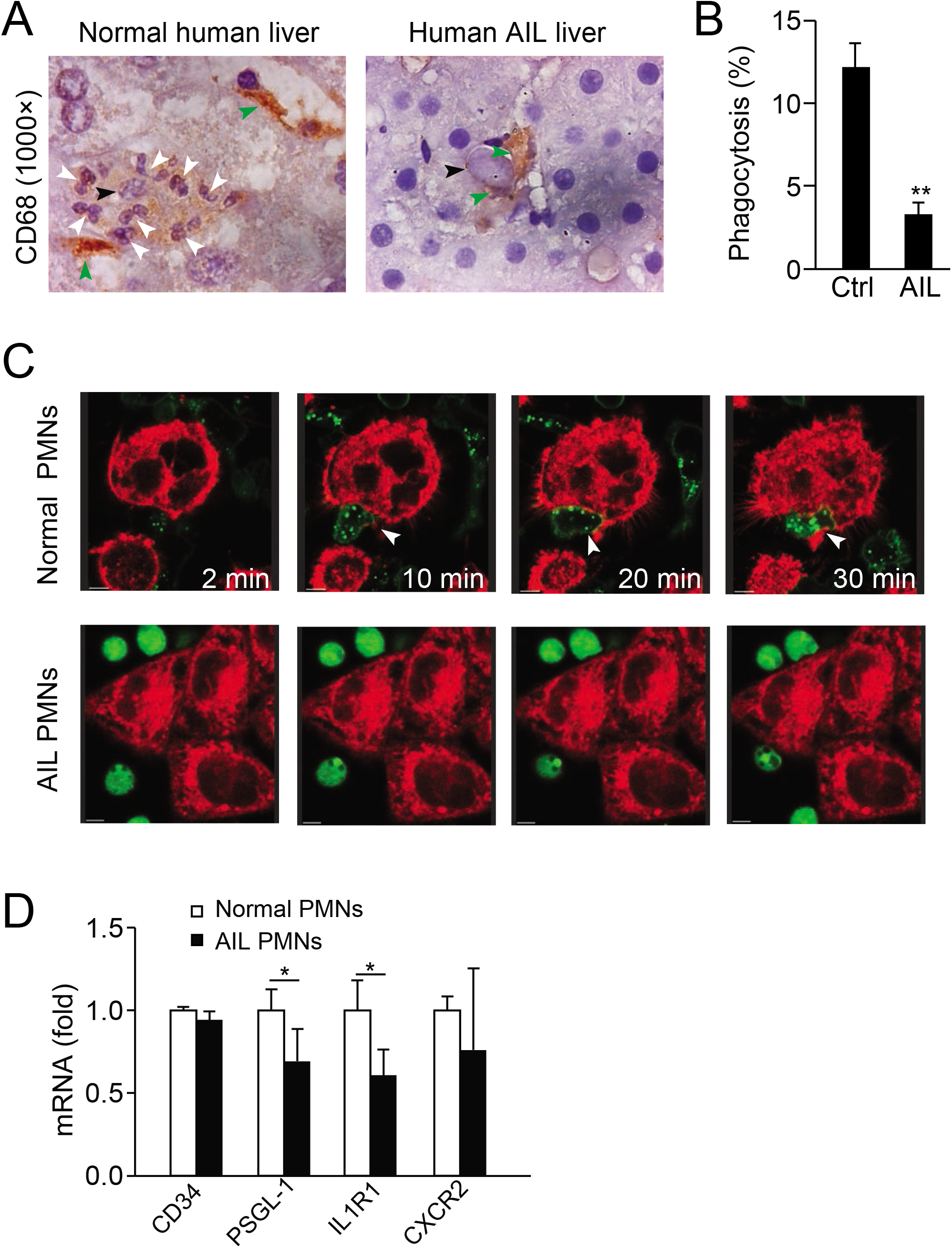
Defective neutrophil perforocytosis in human autoimmune liver disease. (A) Images of CD68 staining from normal or AIL disease human liver samples. Green arrowheads indicate macrophages, black arrowheads indicate apoptotic bodies and white arrowheads point to neutrophils. (B, C) Quantification (B) and images (C) of *in vitro* phagocytosis of apoptotic NCTC cells by neutrophils isolated from normal or AIL disease human patients. White arrowheads point to burrowed neutrophils. Scale bar, 10 μm. *, *P* < 0.05 (Student’s *t*-test). (D) Analysis of CD34, PSLG-1, IL1R1 and CXCR2 expression in neutrophils from normal human and AIL patients by microarray. Data are representative of (A, C) or from three independent experiments (B, D; mean and s.e.m. in B, D).

## Discussion

Our finding of neutrophil burrowing into and clearing of apoptotic hepatocytes under physiological conditions reveals a fundamental mechanism for the removal of effete hepatocytes, helps to solve some mysteries surrounding the apoptotic clearance process and raises several important questions.

### Neutrophils as the new scavengers for apoptotic clearance

Although billions of cells undergo apoptosis daily in our bodies, apoptotic cells are rarely observed in tissues under steady state due to the high efficiency of apoptotic removal by both professional and neighboring nonprofessional phagocytes(Poon et al., 2014). In general, professional phagocytes have much higher phagocytic efficiency and capacity than nonprofessional phagocytes, but they are greatly outnumbered by other cell types in tissues(Elliott and Ravichandran, 2016). Therefore, whether there are unidentified scavengers and/or mechanisms for the prompt removal of dead cells in distinct organs remains unclear. Compared to macrophages or their precursors, monocytes, which make up less than 10% of the total white blood cell population, neutrophils are the most abundant white blood cells (up to 70%) in humans, and thus ideally suited for the swift clearance of apoptotic hepatic cells. As macrophages were induced to phagocytose apoptotic hepatocytes following the depletion of circulating neutrophils, our data show compensatory activation of other phagocytes under these conditions. The basis for this switch, however, is not clear.

Another important question is whether other tissues utilize a similar mechanism. As neutrophils do not phagocytose apoptotic epithelial and endothelial cells, it is possible that distinct signals are specifically released by different types of apoptotic cells. It remains largely unknown whether one or more signals are dominant in each specific tissue and how these signals are orchestrated into a complicated network to swiftly clear dying cells while avoiding tissue inflammation.

### Neutrophil influx without causing tissue damage

Although apoptotic clearance has been considered an immunologically silent process that does not lead to the influx of inflammatory cells or exposure of self-antigens(Poon et al., 2014; Savill et al., 2002), it has also been suggested that this process is not completely immunologically silent (Green et al., 2009). Because neutrophils are proinflammatory cells, they have long been thought to be excluded from apoptotic sites. We observed neutrophil influx into apoptotic hepatic cells (up to 22 neutrophils in one apoptotic hepatocyte), and there was little detectable tissue injury or other inflammatory cells. One reason could be that the neutrophil attraction signals released by apoptotic hepatocytes (e.g., IL-1β and IL-8) are sufficient for attracting neutrophils towards apoptotic cells and inducing subsequent perforocytosis, but do not elicit neutrophil inflammatory functions such as oxidative burst. Therefore, neutrophil recruitment into tissues does not always induce inflammation or tissue damage. In consistent with our observations, neutrophils can express both pro- and anti-inflammatory cytokines(Gideon et al., 2019; Mortaz et al., 2018). In contrast, neutrophil depletion caused defective removal of apoptotic bodies and induced autoantibody generation; thus neutrophils play a vital role in the genesis of AIL disease. Considering that billions of neutrophils patrol tissues under physiological circumstances without causing inflammation(Nicolas-Avila et al., 2017), we concluded that these cells not only provide immune surveillance against infection but also contribute to internal tissue homeostasis, as we report in this study.

### Neutrophil burrowing to maintain tissue integrity

One intriguing question regarding apoptotic clearance is how the tissue maintains integrity while dead cells are being continually removed(Poon et al., 2014). Apoptotic cell detachment from the extracellular matrix by caspase-mediated cleavage or extrusion into the organ lumen has been proposed as the mechanism for maintaining the epithelial barrier during the clearance of dying epithelial cells(Brancolini et al., 1997; Rosenblatt et al., 2001). Our finding that burrowed neutrophils phagocytose apoptotic hepatocytes from the intracellular space provides a novel mechanism for maintaining liver integrity. The neutrophils that phagocytose cells from the inside are efficient at disposing of apoptotic bodies without extruding the cytoplasm and may help to prevent the leakage of toxic bile acids or the release of danger signals that cause tissue damage.

Neutrophil burrowing into other types of cells has also been reported previously in processes not related to apoptotic clearance(Overholtzer and Brugge, 2008). For example, neutrophils can bore into endothelial cells or megakaryocytes to temporarily form so-called cell-in-cell (or emperipolesis, entosis) structures(Overholtzer and Brugge, 2008). The apparent difference is that both the endothelial cells and megakaryocytes entered by neutrophils are viable and nonapoptotic. The purpose of forming cell-in-cell structures with neutrophils and endothelial cells or megakaryocytes is to obtain a passage out of blood vessels or bone marrow, respectively(Overholtzer and Brugge, 2008). It will be interesting to examine whether neutrophils utilize similar invasion mechanisms during these processes.

The important apoptotic clearance function of neutrophils described in this study adds to the repertoire of other known functions of neutrophils(Amulic et al., 2012; Kolaczkowska and Kubes, 2013; Wang et al., 2017). Since the failure to clear dead cells is linked to inflammatory and autoimmune diseases, the critical role of neutrophil-mediated apoptotic clearance may have implications for the pathogenesis and treatment of these diseases.

## Methods

### Cell culture, transfection, and isolation of human and mouse neutrophils

NCTC, HEK-293, U937 cells and HUVECs were cultured in DMEM supplemented with 10% fetal bovine serum and HL60 cells were cultured in RPMI 1640 with 10% fetal bovine serum. For experiments, HL60 cells were differentiated by adding 1.3% DMSO into the medium for 7 days(Xu et al., 2003). To establish the stable RNAi cell lines, we transfected shRNA into HEK293T cells. After virus packaging in HEK293T cells, HL60 cells were infected, screened by puromycin and sorted by flow cytometry(Liu et al., 2012).

For mouse polymorphonuclear neutrophil (PMN) isolation from bone marrow(Wang et al., 2016), mice were euthanized, and the femurs and tibias were removed and flushed by a 27-gauge needle with a 10-mL syringe filled with calcium and magnesium-free Hank’s balanced salt solution (HBSS) plus 0.1% BSA. Cells were then centrifuged and resuspended in HBSS. After being filtered through a 40-μm strainer, cells in 3 mL of HBSS were loaded onto a pre-prepared gradient solution (3 mL of NycoPrep on top and 3 mL of 72% Percoll on bottom). The samples were centrifuged at 2,400 rpm at room temperature for 20 min with the brake off. The middle layer was collected and washed once in HBSS. Sterilized distilled water (9 mL) was added for 22 s and thereafter 1 mL of 10× PBS was added to remove red blood cells. Finally, the cells were collected and resuspended in HBSS or medium (neutrophil purity >90%). All animal experiments were performed under the protocol approved by The Institutional Animal Care and Use Committee of the State Key Laboratory of Biotherapy, West China Hospital, Sichuan University.

For the isolation of human neutrophils (Liu et al., 2015), blood was collected from healthy human donors or those with AIL disease. Erythrocytes were removed using dextran sedimentation (4.5% dextran) followed by hypotonic lysis using distilled water. Neutrophils were isolated from the resulting cell suspension using discontinuous Percoll gradient centrifugation. This procedure yielded >95% neutrophil purity and >95% viability as assessed by flow cytometry.

For the isolation of human primary hepatocytes, liver tissues from a fresh liver hemangioma surgery were suspended in DPBS and washed for 5-10 times. Surrounding connective tissues and adipose tissues were removed with a surgery knife and remaining liver tissues were further sliced into small fragments (around 1 mm^2^), then washed with DPBS for three times and centrifuged at 800 rpm for 3 min. Tissue samples were resuspended in 5 times volume of collagenase IV medium and incubated at 37℃ for 30 min. After collagenase digestion, samples were centrifuged at 1000 rpm for 5min, and resuspended in RPMI 1640 with antibiotics to precipitate cells, then centrifuged at 1500 rpm for 5 min. Isolated cells were cultured in RPMI 1640 medium with insulin (1: 1000), glucagon (1:1000), glucocorticoid (1:1000), HGF (10 ng/mL), EGF (10ng/mL). Studies using human primary cells and samples were approved by the Institutional Review Board of West China Hospital, Sichuan University and Fudan University.

### Intravital microscopy, immunostaining of liver sections, 3D image reconstruction and electron microscopy

For intravital microscopy(Marques et al., 2015), freshly prepared anti-Ly6G 1A8 FITC, anti-F4/80 PE and Annexin V pacific blue were injected intravenously into mice 1 h before imaging. After anesthesia, mice were attached to a surgical stage and subjected to midline laparotomy, and all vessels in the skin were cauterized. The falciform ligament between the liver and diaphragm was subsequently cut by holding the xiphoid with a knot made of suture thread. Mice were then moved to the imaging stage, and the right lobe was exposed. The ventral side of the mouse was placed on a Plexiglas stage attached to the microscope stage of a single photon microscope (NiKON A1RMP) or Cellvizio System for the imaging of apoptotic cells (labeled with Annexin V), Kupffer cells (anti-F4/80) and neutrophils (anti-Ly-6G 1A8). Microscope acquisition settings were as follows: 405-nm laser power, 15.4%, PMT high voltage (HV), 115; 488-nm laser power, 27.9%, PMT HV, 76; 543-nm laser power, 41.4%, PMT HV, 108; pinhole size, 40 μm; 20× objective.

Neutrophils burrowing inside apoptotic hepatocytes were analyzed by IMARIS software (adopted from the analysis of vesicles in cells). Annexin V blue was set as the source channel to detect apoptotic hepatocyte boundary, and anti-Ly6G green was set as the source channel to detect neutrophils inside blue cells. Only neutrophils inside hepatic cells can be detected. The cells objects were calculated and displayed. The cell positions of both neutrophils and hepatocytes, and the distances from neutrophils to the hepatocyte border were recorded.

For immunostaining of liver sections, fresh mouse liver samples were fixed in freshly prepared 4% PFA and cryoprotected in 30% sucrose/PBS solution. Thick cryosections (45 μm) were cut and washed three times with PBS. Tissue sections were incubated in blocking buffer (10% donkey serum and 0.2% Triton X-100 in PBS) for 1 h at room temperature and then in a primary antibody solution (5% donkey serum and 0.2% Triton X-100 in PBS; rat anti-Ly-6G antibody, 1:500; goat anti-E-cadherin antibody, 1:200) overnight at 4°C. Next, the sections were washed three times in PBS, incubated for 1 h in the appropriate secondary antibody solution containing DAPI for 2 h at 4°C, and then washed three times in PBS. The immunostained sections were mounted onto slides with an aqueous mounting medium. All tissue sections were scanned by a Zeiss 710 confocal microscope. The Z-stack scanned images were edited by Zen 2009 software.

To observe the liver ultrastructure with TEM, mouse livers were removed, fixed (2.5% glutaraldehyde in 0.1 M cacodylate buffer, pH 7.4), and cut into multiple 1 mm × 1 mm strips in fluid; several strips were simultaneously cut into 1 mm × 1 mm × 1 mm blocks for TEM. Blocks were transferred to wash buffer (0.1 M cacodylate buffer, pH 7.4) to remove glutaraldehyde and postfixed for 1 h in 1% osmium tetroxide. The specimens were dehydrated in a graded series of ethanol (75%, 85%, 95%, and 100%) and embedded in Epon812. Ultrathin sections were obtained and then observed with a HITACHI H600 TEM. Apoptotic liver cells were determined based on changes in nuclear morphology (chromatin condensation and fragmentation). Neutrophils were determined by polymorphonuclear morphology.

### Neutrophil depletion and macrophage depletion in mice

Two methods were used to deplete neutrophils in mice. For antibody-based depletion, wild-type mice were injected i.p. with anti-mouse Ly6G clone 1A8 (Bio X Cell) supernatant containing 0.5 mg protein every 48 h for 1 month. Control animals were injected with isotype-matched normal control antibody. For genetic neutrophil ablation, diphtheria toxin receptor (DTR)-expressing Rosa26iDTR mice (Buch et al., 2005; Saito et al., 2001) (The Jackson Laboratory) were bred onto the *Mrp8*-Cre genetic background (The Jackson Laboratory). Cre expression in MRP-expressing cells, mainly granulocytes, will activate DTR expression in only MRP-expressing cells. At the age of 3 months, the Rosa26iDTR/Mrp8-Cre mice and WT control littermates (Rosa26iDTR) were treated with diphtheria toxin (DT, 20 ng/g BW, i.p., daily for 3 days) to induce Mrp8^+^ cell depletion.

Clodronate-liposome (YEASEN, 40337ES05) was used to deplete mouse macrophage. WT mice were injected i.v. with 100 μl of clodronate-liposome per 10 g of body weight. 24 h later, mice were injected i.v. with 100 μl of clodronate-liposome per 10 g of body weight again. The operation was repeated every 72 h after the second injection for 1 month. Neutrophil and macrophage depletion were confirmed by neutrophil and macrophage count in all experimental animals.

### Apoptosis assay

Apoptosis of NCTC cells was induced by puromycin(Chow et al., 1995). A total of 1 × 10^6^ NCTC cells were plated in a 35 mm glass-bottom dish (MatTek) for 10 h, and then treated with puromycin (2.5 μg/ml) for 12 h. Apoptosis was assayed with an Alexa Fluor 488 Annexin V/Dead Cell Apoptosis Kit (Thermo Fisher) and analyzed by flow cytometry to measure the fluorescence emission at 530 nm and 575 nm (or equivalent) using 488 nm excitation.

### Fluorescent labeling of cells

The fluorescent membrane stain PKH-26 (red) (Sigma-Aldrich) was used to label nonapoptotic and apoptotic NCTC and HEK293 cells and HUVECs. PKH-67 (green) was used to label HL60 cells and human neutrophils. Cells (2 × 10^6^) were washed in serum-free DMEM or RPMI 1640 medium, and the cell pellet was resuspended in PKH-26 or PKH-67 staining solution and incubated for 5 min at room temperature. After stopping the labeling reaction by adding an equal volume of serum, the cells were washed three times with complete medium.

### *In vitro* phagocytosis assay

Phagocytosis was quantified by flow cytometry using a pH sensitive dye, PHrodo-red (weak fluorescent at neutral pH but high fluorescent with a drop in pH) to measure engulfment. 1 × 10^6^ human neutrophils or differentiated HL60 cells were labelled with CFSE green and then incubated with apoptotic or nonapoptotic NCTC cells (labeled with PHrodo-red, 10 μL PHrodo-red dye mixed with 100 μL PowerLoad concentrate, and diluted into 10 mL of HBSS, PH 7.4 for 1 h). The cells were then washed with HBSS and analyzed by flow cytometry. The forward scatter (FCS) reflects cell size, whereas the sideward scatter (SSC) reveals the degree of granularity of the cell. Apoptotic NCTC cells with burrowed neutrophils increased in cell size and granularity, causing increased diffraction of the laser beam, revealed by spreading of the dots to the upper right corner. The population of double fluorescent HL60 cells in this subgroup was calculated, and these cells were considered phagocytosing cells.

### Cytokine screening and microarray

Cytokines secreted from nonapoptotic and apoptotic NCTC cells were screened with the Human Inflammatory Cytokines Multi-Analyte ELISArray Kit (QIAGEN). Nonapoptotic and apoptotic NCTC cell supernatants were prepared in replicate serial dilutions of the Antigen Standard. Cytokine secretion was further determined with the ELISA kit.

For microarray, total RNA from normal human or AIL patient neutrophils was isolated using RNeasy Total RNA Isolation Kit (Qiagen, GmBH, Germany)/ TRIzol reagent (Life technologies, Carlsbad, CA, US) according to the manufacturer’s instructions, and purified by using RNeasy Mini Kit (Qiagen, GmBH, Germany). Total RNA was checked for a RIN number to inspect RNA integration by an Agilent Bioanalyzer 2100 (Agilent technologies, Santa Clara, CA, US). RNA samples of each group were then used to generate biotinylated cRNA targets for the Agilent SurePrint Gene Expression Microarray. The biotinylated cRNA targets were then hybridized with the slides. After hybridization, slides were scanned on the Agilent Microarray Scanner (Agilent technologies, Santa Clara, CA, US). Data were extracted with Feature Extraction software 10.7 (Agilent technologies, Santa Clara, CA, US). Raw data were normalized by Quantile algorithm, R package “limma”. Heatmap plots were done by a R package “pheatmap” of the target genes. GO\pathway enrichment analysis was done use Fisher’s exact test by R package “clusterProfiler” of the target genes. GO categories\Pathway with Fisher’s exact test P values < 0.05 were selected.

### Statistical analysis

All experiments were performed at least three times. Representative data are shown in the paper. An independent experiments two-tailed Student’s *t*-test was used as the statistical assay for comparisons. Significant differences between samples were indicated by *P* < 0.05.

## Author contributions

Conceptualization: LC, HS, JZ, ABM, JX

Methodology: LC, HS, JZ, JL, XF, WL, YQ, YT, SP, LY, LZ, LM, LL, XH, YJ, YH, YZ, XZ, YZ, YZ, YW

Investigation: LC, HS, JZ, JL, XF, WL, YQ, YT, SP, LY, LZ, LM, LL, XH, YJ, YH, YZ, XZ, YZ, JX

Visualization: LC, HS, JZ, JL, XF, WL, YQ, YT, SP, LY, LZ, LM, LL, XH, YJ, YH, YZ, XZ, YZ

Funding acquisition: LZ, YZ, YZ, YW, ABM and JX Project administration: LC, HS, JZ, JX

Supervision: LC, HS, JX Writing: LC, HS, ABM, JX

## Acknowledgement

We thank Yu Fu and Ruixue Li (both from Guangzhou Regenerative Medicine and Health Guangdong Laboratory Imaging Center) for helpful imaging service. This work was supported in part by NSFC grant 32170750 and SSTP grant 19YYJC2572.

---

**Fig. S1.**
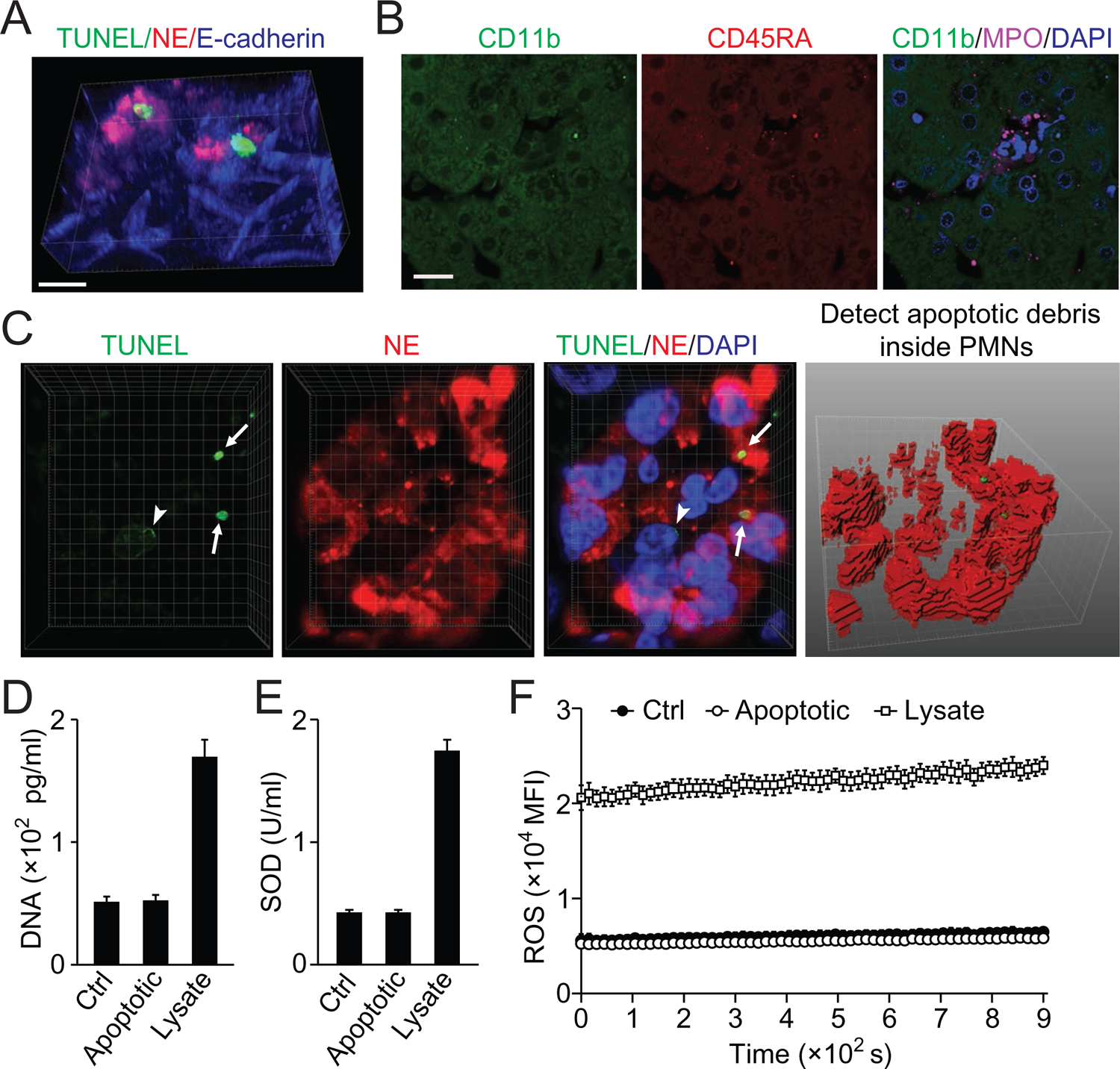
Little leakage during perferocytosis. (A) Reconstructed 3D images of neutrophils (NE staining, red) burrowed into apoptotic hepatocytes (TUNEL, green). (B) Few CD11b^+^ or CD45RA^+^ cells around the perferocytosis site. Bar, 20μm. (C) Fluorescent images of human liver tissue with neutrophils scavenging apoptotic liver cells. TUNEL staining (green) shows apoptotic nucleus (white arrowheads) and apoptotic nuclear debris (white arrows). The apoptotic debris is phagocytosed by burrowed neutrophils (labeled with NE-red, analyzed and projected with IMARIS software). The burrowing neutrophils have intact nucleus (shown by DAPI staining) with negative TUNEL signals. (D-F) Extracellular levels of DNA (D), SOD (E), and ROS (F) are not significantly changed during phagocytosis of apoptotic NCTC cells by HL60 cells as compared with that of nonapoptotic control NCTC cells. Cell lysate was used as positive control. Data are representative of (A-C) or from (D-F) 3 independent experiments (mean and s.e.m. in D-F).

**Fig. S2.**
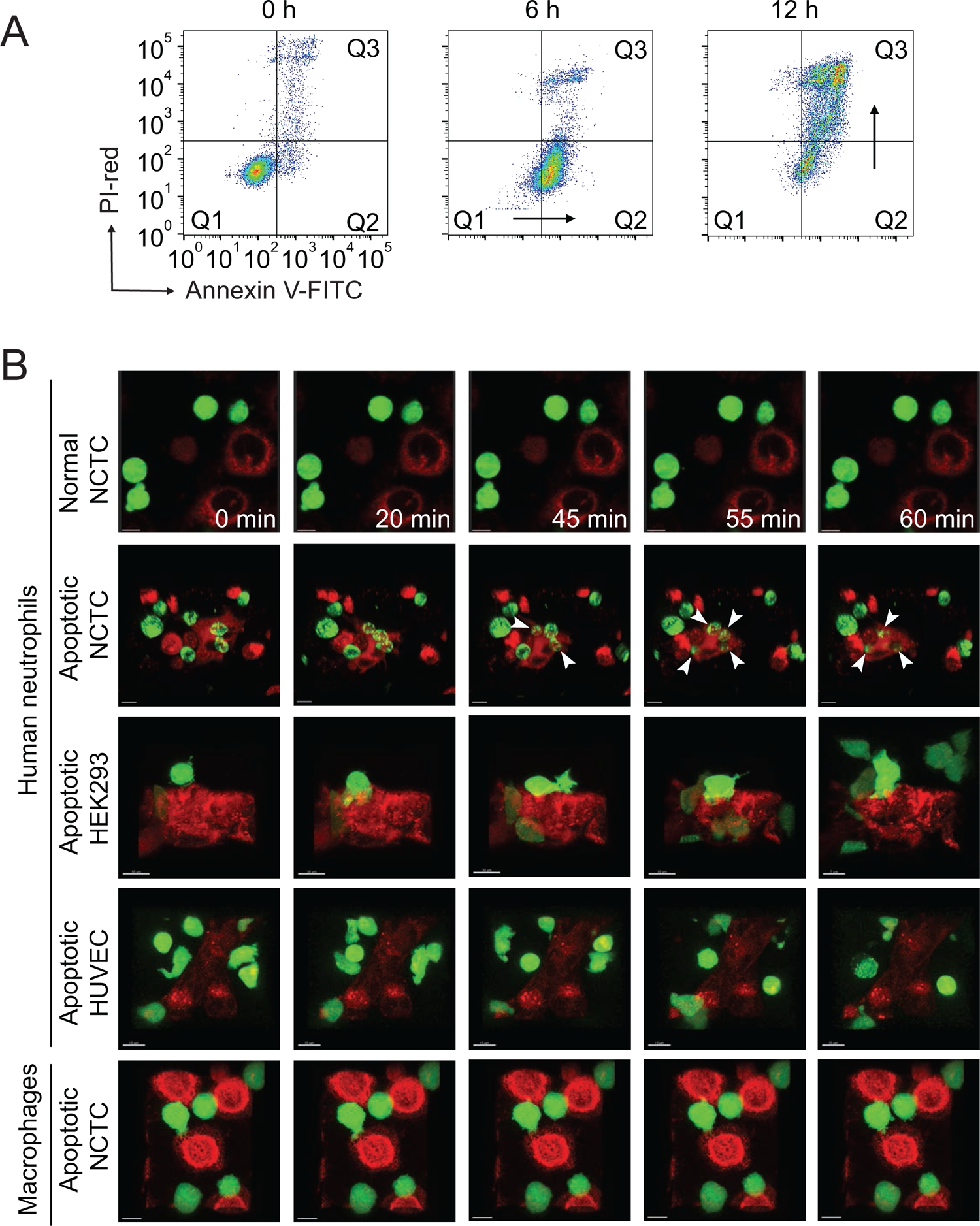
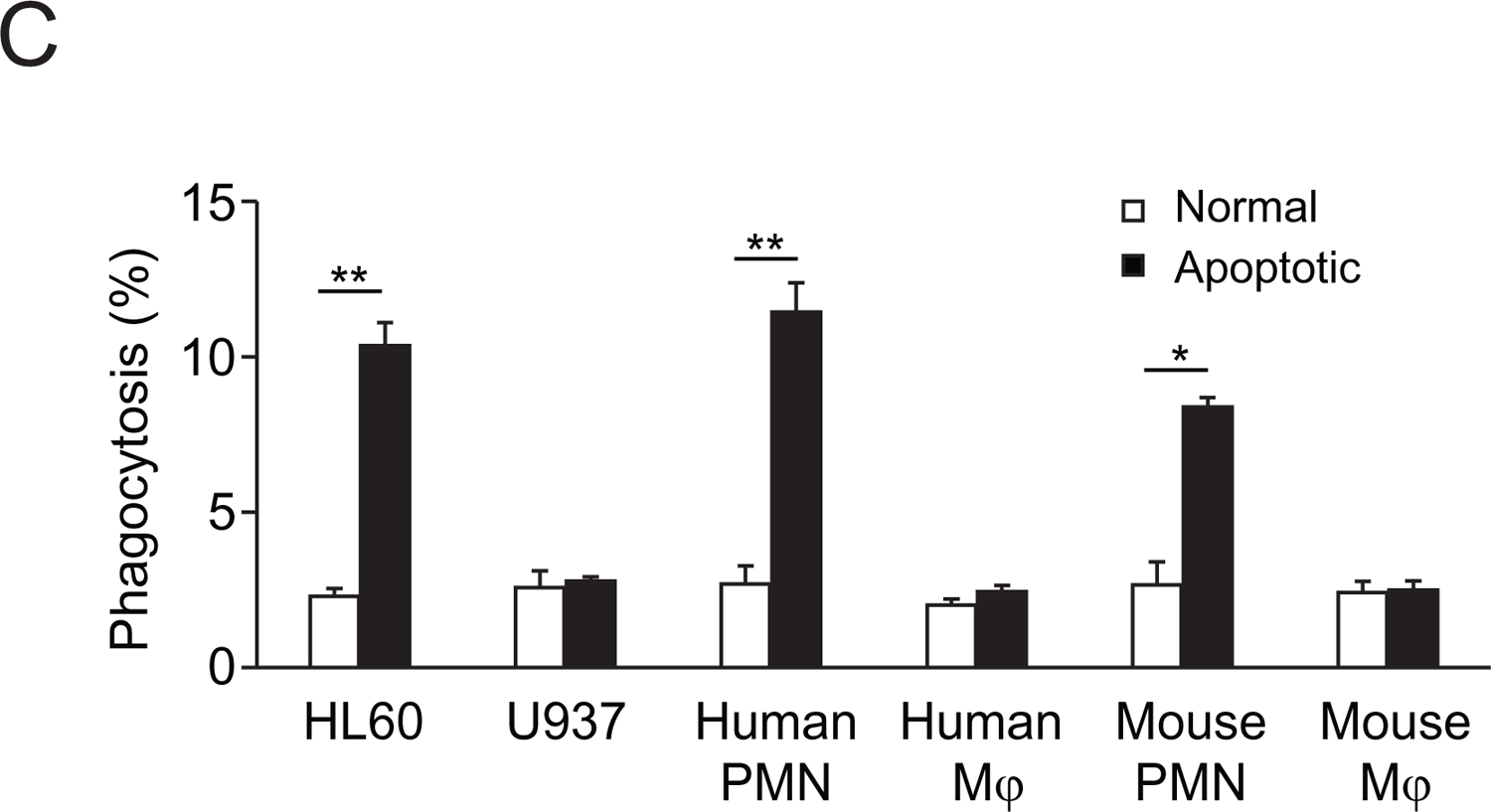
Neutrophils selectively phagocytose apoptotic liver cells. (A) Flow analysis of NCTC cells treated with puromycin at indicated time points in the presence of PI-red and Annexin-V-FITC. A lag time seen between PtdSer^+^ and PI^+^ is a characteristic feature of apoptotic cells, the cell population moves first from Q1 (AnnexinV^-^/PI^-^) to Q2 (AnnexinV^+^/PI^-^), then moves to Q3 (AnnexinV^+^/PI^+^. In contrast, necrotic cells immediately move from Q1 to Q3 without passing though the Q2 stage. (B) PKH67 (green)-labeled human neutrophils or macrophages interacting with PKH26 (red)-labeled NCTC, HEK293 cells and HUVECs at the indicated times. Neutrophils only burrowed inside apoptotic NCTC cells (second row, white arrowheads point to the burrowing neutrophil). Bar, 10 µm. (C) Phagocytosis of HL60, U937, human/mouse neutrophils (PMN), or macrophages (Mj), towards nonapoptotic and apoptotic NCTC cells. *, *p* < 0.05, **, *p* < 0.01, (Student’s *t*-test). Data are representative of (A, B) or from (C) three independent experiments (mean and s.e.m. in C).

**Fig. S3.**
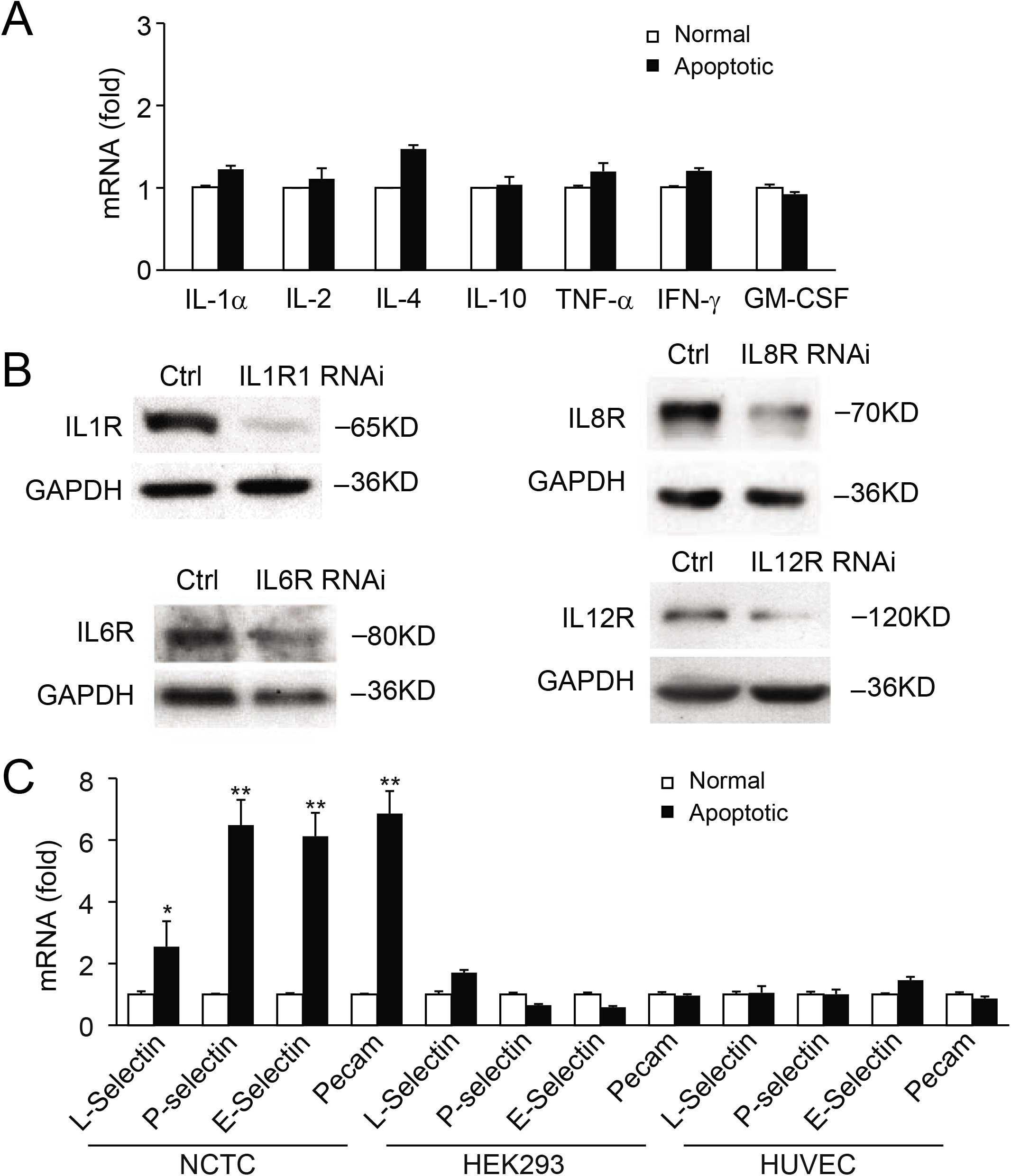
Cytokines and cell surface receptors for perferocytosis. (A) Cytokines secreted by normal nonapoptotic and apoptotic NCTC cells are not significantly changed, includig IL-1α, IL-2, IL-4, IL-10, TNF-α, IFN-γ, and GM-CSF. (B) Immunblots of target proteins (IL-1b, IL-8, IL-6, IL-12 receptors) in non-treated (Ctrl) and RNAi-treated HL60 cells. RNAi knockdown efficiency is ranged from 70-90%. Data are from (A) or representative of (B) three independent experiments (mean and s.e.m. in A).

**Fig. S4.**
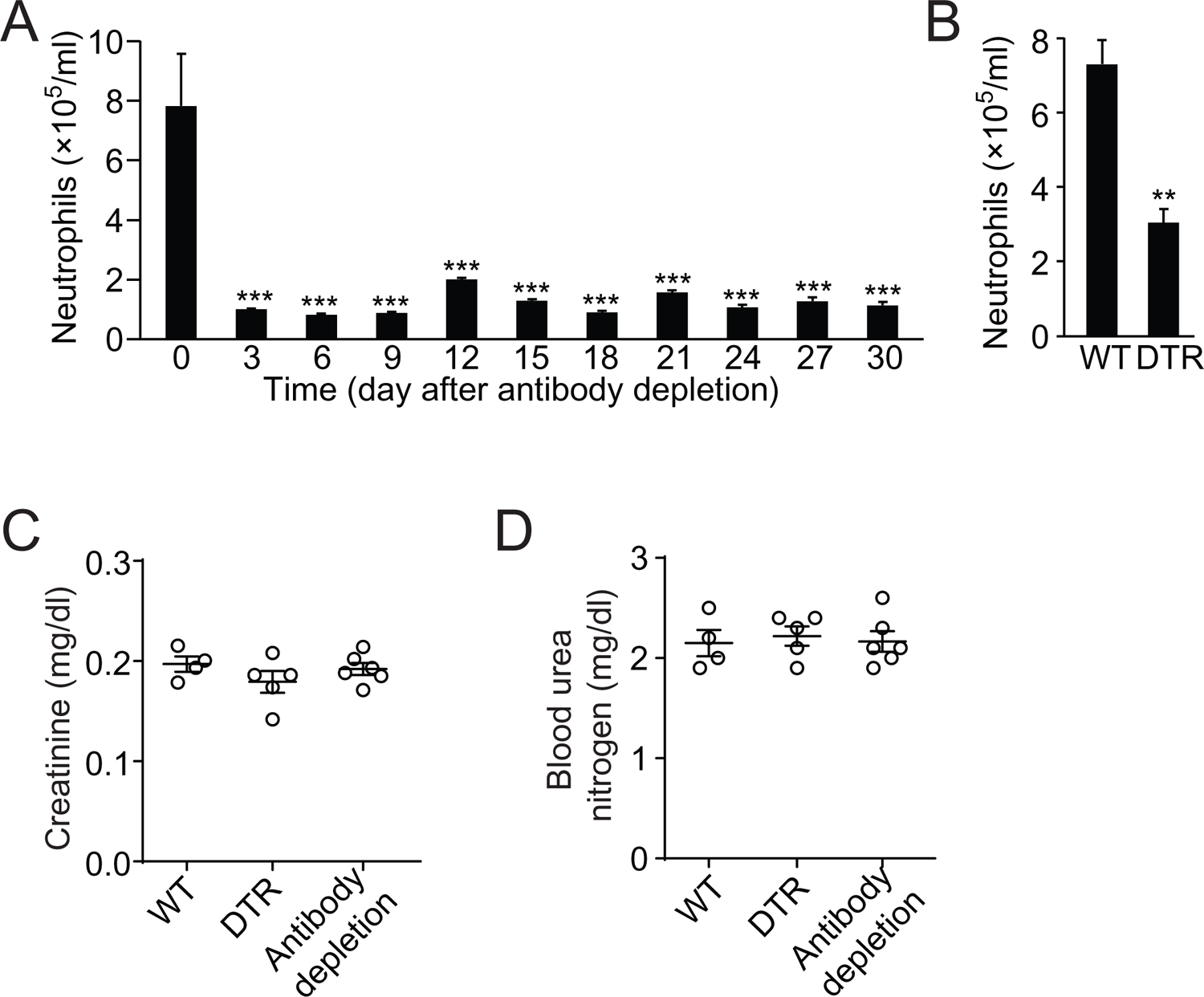
Neurophil depletion impairs liver functions. (A,B) Blood neutrophil count after antibody depletion at times indicated (A) or in WT and MRP8cre/DTR mice (B). (C,D) Analysis of creatinine (C) and blood urea nitrogen (D) in WT, MRP8Cre/DTR, and neutrophil depleted (by antibody) mice. Data are from 3 independent experiments (mean and s.e.m. in A-D).

**Table 1.** The cell numbers of burrowed neutrophils (PMNs) in apoptotic hepatocytes from human liver tissues. 2D images were taken from regular tissue sections (5 μm thick), and 3D images were taken from 45 μm thick tissue sections and reconstructed by confocal microscope.

| Perferocytosis stage |  | I | II | III |
| --- | --- | --- | --- | --- |
| 2D images | Apoptotic hepatocytes (total 241) | 37 | 130 | 74 |
|  | Burrowing PMNs (total 1716) | 312 | 931 | 473 |
| | PMNs per apoptotic hepatocyte (mean ± sem) | 8.4 ± 4.2 | 7.2 ± 3.4 | 6.4 ± 3.4 |
| 3D images | Apoptotic hepatocytes (total 60) | 28 | 18 | 14 |
|  | Burrowing PMNs (total 820) | 406 | 348 | 66 |
| | PMNs per apoptotic hepatocyte (mean ± sem) | 14.5 ± 10.0 | 19.3 ± 15.5 | 4.7 ± 2.4 |

**Table 2.**
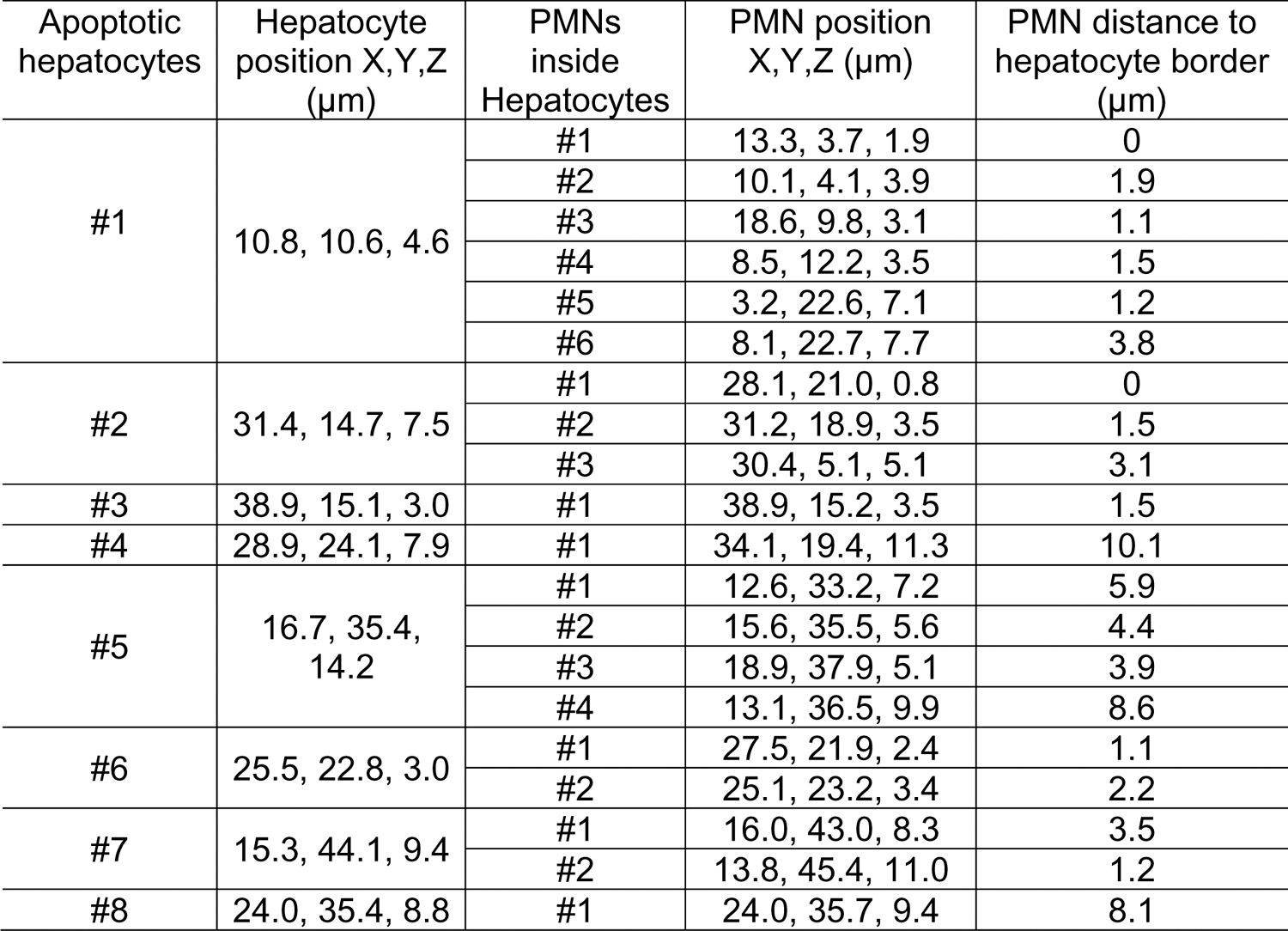
Analysis of neutrophils burrowing inside apoptotic hepatocytes from human liver samples. Reconstructed 3D images of neutrophils inside apoptotic hepatocytes (as shown in Figure 1D) are remodeled and analyzed with IMARIS software: anti-E-Cadherin blue staining is set as the source channel to detect hepatic cell boundary, and anti-NE red staining is set as the source channel to detect neutrophils inside blue cells. Only neutrophils inside hepatic cells can be detected. The cell positions of both neutrophils and hepatocytes, and the distances from neutrophils to the hepatocyte border are recorded below (a total of 8 apoptotic hepatocytes are analyzed).

**Table 3.** CD68^+^ cells’ distribution in human liver tissues with or without apoptotic hepatocytes (perferocytosed by PMNs). 5 μm thick tissue sections were used to stain and count CD68^+^ cells.

| | Total area<br>($\mu\text{m}^2$) | CD68<sup>+</sup> cells | Apoptotic<br>hepatocytes | CD68<sup>+</sup> cells/$10^4 \mu\text{m}^2$<br>(mean ± sem) |
| --- | --- | --- | --- | --- |
| Liver tissue<br>with PMN<br>perferocytosis | 579175 | 280 | 40 | 5.8 ± 2.2* |
| Liver tissue<br>without PMN<br>perferocytosis | 403025 | 267 | 0 | 6.7 ± 2.8* |

\*, $P = 0.0962$ (student's $t$ test), CD68<sup>+</sup> cell numbers are not different in liver tissues with or without PMN perferocytosis.

**Table 4.**
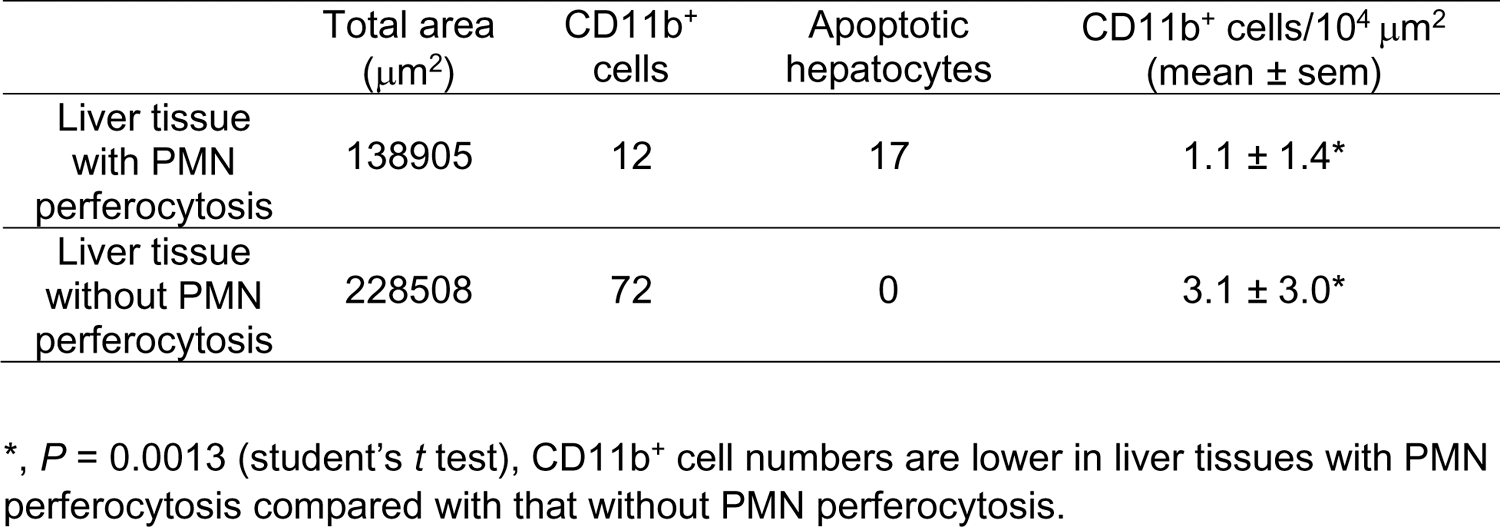
The distribution of CD11b^+^ cells in human liver tissues with or without apoptotic hepatocytes (perferocytosed by PMNs). 45 μm thick tissue sections were used to stain and count CD11b^+^ cells.

**Table 5.** The distribution of CD45RA^+^ cells in human liver tissues with or without apoptotic hepatocytes (perferocytosed by PMNs). 45 μm thick tissue sections were used to stain and count CD45RA^+^ cells.

| | Total area<br>($\mu\text{m}^2$) | CD45RA<sup>+</sup><br>cells | Apoptotic<br>hepatocytes | CD45RA<sup>+</sup> cells/$10^4 \mu\text{m}^2$<br>(mean ± sem) |
| --- | --- | --- | --- | --- |
| Liver tissue<br>with PMN<br>perferocytosis | 57367 | 7 | 10 | 1.4 ± 1.2* |
| Liver tissue<br>without PMN<br>perferocytosis | 280880 | 54 | 0 | 1.9 ± 1.9* |

\*, $P = 0.309$ (student's $t$ test), CD45RA<sup>+</sup> cell numbers are not different in liver tissues with or without PMN perferocytosis.

**Table 6.**
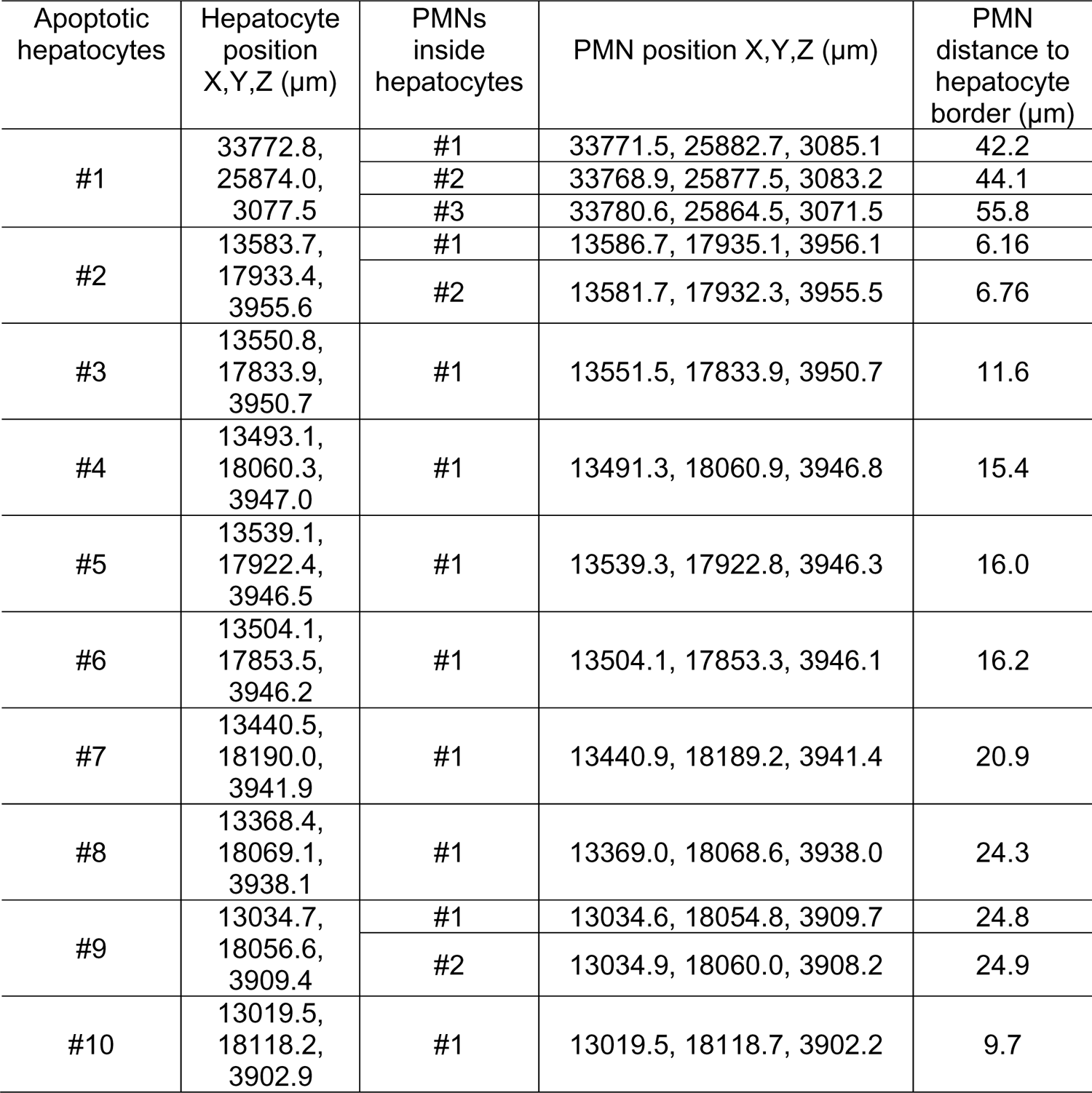
Analysis of neutrophils burrowing inside apoptotic hepatocytes from mouse liver samples. Intravital images of neutrophils inside apoptotic hepatocytes (as shown in Figure 2A) are remodeled and analyzed with IMARIS software: Annexin V blue staining is set as the source channel to detect hepatic cell boundary, and Ly-6G green staining is set as the source channel to detect neutrophils inside blue cells. Only neutrophils inside hepatic cells are detected and calculated. The cell positions of both neutrophils and hepatocytes, and the distances from neutrophils to the hepatocyte border are recorded below (a total of 10 apoptotic hepatocytes are analyzed).

| Apoptotic hepatocytes | Hepatocyte position X,Y,Z (μm) | PMNs inside hepatocytes | PMN position X,Y,Z (μm) | PMN distance to hepatocyte border (μm) |
| --- | --- | --- | --- | --- |
| #1 | 33772.8, 25874.0, 3077.5 | #1 | 33771.5, 25882.7, 3085.1 | 42.2 |
|  |  | #2 | 33768.9, 25877.5, 3083.2 | 44.1 |
|  |  | #3 | 33780.6, 25864.5, 3071.5 | 55.8 |
| #2 | 13583.7, 17933.4, 3955.6 | #1 | 13586.7, 17935.1, 3956.1 | 6.16 |
|  |  | #2 | 13581.7, 17932.3, 3955.5 | 6.76 |
| #3 | 13550.8, 17833.9, 3950.7 | #1 | 13551.5, 17833.9, 3950.7 | 11.6 |
| #4 | 13493.1, 18060.3, 3947.0 | #1 | 13491.3, 18060.9, 3946.8 | 15.4 |
| #5 | 13539.1, 17922.4, 3946.5 | #1 | 13539.3, 17922.8, 3946.3 | 16.0 |
| #6 | 13504.1, 17853.5, 3946.2 | #1 | 13504.1, 17853.3, 3946.1 | 16.2 |
| #7 | 13440.5, 18190.0, 3941.9 | #1 | 13440.9, 18189.2, 3941.4 | 20.9 |
| #8 | 13368.4, 18069.1, 3938.1 | #1 | 13369.0, 18068.6, 3938.0 | 24.3 |
| #9 | 13034.7, 18056.6, 3909.4 | #1 | 13034.6, 18054.8, 3909.7 | 24.8 |
|  |  | #2 | 13034.9, 18060.0, 3908.2 | 24.9 |
| #10 | 13019.5, 18118.2, 3902.9 | #1 | 13019.5, 18118.7, 3902.2 | 9.7 |

**Table 7.** Apoptotic hepatocytes burrowed with PMNs or monocytes in normal human or AIL liver samples. 45 μm thick tissue sections were used to stain and count apoptotic hepatocytes.

|  | Total liver<br>sample<br>numbers | Apoptotic<br>hepatocytes | Apoptotic<br>hepatocytes<br>burrowed by<br>PMNs | Apoptotic<br>hepatocytes<br>burrowed by<br>monocytes |
| --- | --- | --- | --- | --- |
| Normal liver<br>samples | 32 | 227 | 227 | 0 |
| AIL liver<br>samples | 22 | 110 | 8 | 40 |

**Video S1, S2: Mouse neutrophils burrowed and digested apoptotic hepatocytes in the mouse liver.** Neutrophils were labeled with an i.v. injection of anti-Ly6G antibody (green) and apoptotic cells were labeled with Annexin V (red). Time-lapse images were acquired by the Cellvizio system. The cytoplasm of apoptotic hepatocyte with neutrophil burrowing quickly dwindled and finally disappeared. 2 out of 13 apoptotic hepatocytes are shown in video S1 and S2.

**Video S3: Human neutrophils burrowed inside apoptotic hepatocytes**. A human neutrophil (green, pointed by white arrow) was burrowing into an apoptotic hepatocyte (red). Time-lapse video was recorded for 60 minutes and followed by reconstructed 3D images at 20 and 60 min. At 20 min, the neutrophil partially entered the hepatic cell and did not phagocytose (neutrophil was extracted and analyzed by IMARIS software and no red dye inside the neutrophil). At 60 min, the neutrophil completely burrowed inside the hepatocyte and started to phagocytose hepatocyte (with uptake of red dye).

